# A genetic switch for male UV-iridescence in an incipient species pair of sulphur butterflies

**DOI:** 10.1101/2021.05.21.445125

**Authors:** Vincent Ficarrotta, Joseph J. Hanly, Ling S. Loh, Caroline M. Francescutti, Anna Ren, Kalle Tunström, Christopher W. Wheat, Adam H. Porter, Brian A. Counterman, Arnaud Martin

## Abstract

Mating cues evolve rapidly and can contribute to species formation and maintenance. However, little is known about how sexual signals diverge and how this variation integrates with other barrier loci to shape the genomic landscape of reproductive isolation. Here, we elucidate the genetic basis of UV iridescence, a courtship signal that differentiates the males of *Colias eurytheme* butterflies from a sister species, allowing females to avoid costly heterospecific matings. Anthropogenic range expansion of the two incipient species established a large zone of secondary contact across the eastern US with strong signatures of genomic admixtures spanning all autosomes. In contrast, Z chromosomes are highly differentiated between the two species, supporting a disproportionate role of sex chromosomes in speciation known as the large-X (or large-Z) effect. Within this chromosome-wide reproductive barrier, linkage mapping indicates that *cis-*regulatory variation of *bric a brac* (*bab*) underlies the male UV-iridescence polymorphism between the two species. Bab is expressed in all non-UV scales, and butterflies of either species or sex acquire widespread ectopic iridescence following its CRISPR knock-out, demonstrating that Bab functions as a suppressor of UV-scale differentiation that potentiates mating cue divergence. These results highlight how a genetic switch can regulate a premating signal and integrate with other reproductive barriers during intermediate phases of speciation.

**Significance statement:** Incipient species are at an intermediate stage of speciation where reproductive isolation is counteracted by the homogenizing effects of gene flow. Human activity sometimes leads such species to reunite, as seen in the Orange Sulphur butterfly, which forms large hybridizing populations with the Clouded Sulphur in alfalfa fields. Here we show that the sex chromosome maintains these species as distinct, while the rest of their genome is admixed. Sex chromosomes notably determine which males display to females a bright, iridescent ultraviolet signal on their wings. Genetic mapping, antibody stainings, and CRISPR knock-outs collectively indicate that the gene *bric a brac* controls whether UV-iridescent nanostructures develop in each species, elucidating how a master switch gene modulates a male courtship signal.

## Introduction

Premating signals such as pheromones, calls and displays often differ between sexes and species, and by helping animals to tell one another apart, they are integral to the formation of reproductive barriers during speciation itself (1, 2). Mating factors can diverge early in the speciation process due to local adaptation or later due to sexual selection that prevents the generation of unfit hybrids (3). While a coupling of premating and postmating isolation mechanisms is thought to be required for the completion of speciation (4), how mating cue variation actually coincides with other barrier loci to split lineages remains elusive in the empirical literature (5–7).

Previous work on the genetics of hybridization between the sulphur butterflies *Colias eurytheme* and *Colias philodice* highlights their potential for the study of intermediate phases of speciation with gene flow. Initially restricted to the Western US, the range of *C. eurytheme* expanded following both the spread of agricultural alfalfa and the reduction in forest cover in the past 200 years into regions once limited to *C. philodice* (8). As a result, the two species occur in secondary sympatry throughout an anthropogenic contact zone that includes the eastern United States and southern Canada. Both pre- and postzygotic reproductive barriers maintain species status in this system. However, heterospecific matings happen at increased frequency in dense populations (9, 10), partly because males can locate newly emerged females incapable of performing mate rejection behaviours [teneral mating (11)]. Hybrid female sterility forms an intrinsic postzygotic barrier that affects one of the two heterospecific crosses: oogenesis fails in female offspring that inherited a *C. eurytheme* W chromosome and a *C. philodice* Z chromosome (12, 13). This incompatibility is sex-linked and implies that to produce fully fertile progeny, *C. eurytheme* females must select males that are homozygous for a conspecific Z chromosome. An iridescent ultraviolet (UV) pattern acts as a visual mating cue in males and accurately displays their Z-chromosome status to females (9, 14) (**Fig. 1A and Fig. S1**). UV occurs on the dorsal wing surfaces of *C. eurytheme* males only. The Mendelian *U*-locus controls this interspecific variation and was previously mapped to the Z chromosome (15): *C. eurytheme* homozygous recessive males are UV-iridescent (*u/u*), advertising two compatible Z chromosomes to *C. eurytheme* females. Incompatible mates such as *C. philodice* males (*U/U*) and heterozygous hybrids (*U/u*) bear the dominant allele and lack UV. Finally, the female preference trait itself is also linked to the Z chromosome (14). This Z-linked inheritance of genetic incompatibility, mating signal and mating preference supports an “indicator” model of speciation, which was previously theorized as a system where the mating cue can signal species identity and enable selection against hybrids (3, 16),. In this study, we examine the genomic footprint of sex-linked reproductive barriers, and fine-map the allelic variation that switches on the male UV signal in *C. eurytheme*.

**Fig. 1.**
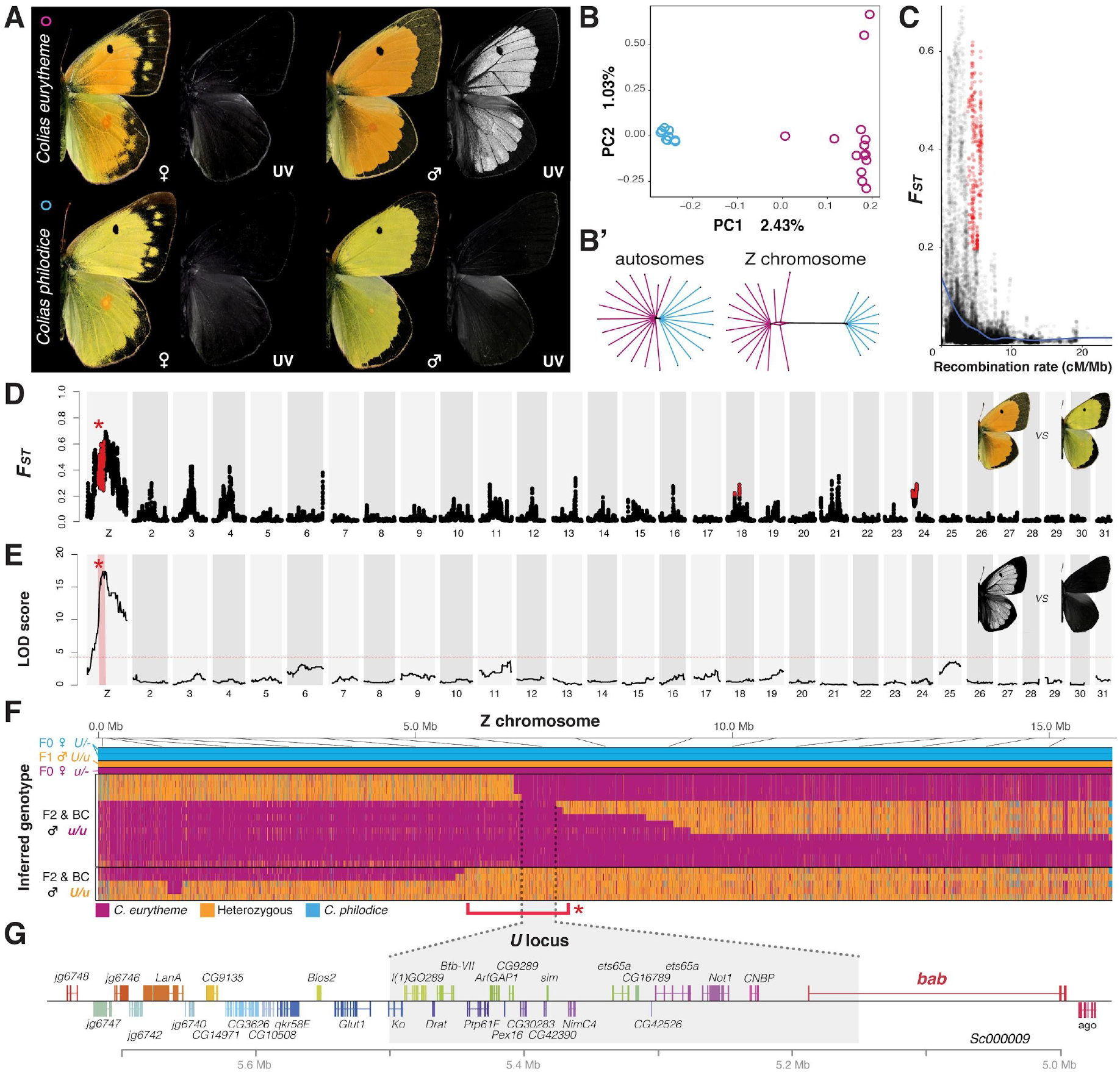
Large-Z architecture of species differentiation includes the *U* locus candidate gene *bab*. (**A**) UV-iridescence differentiates males from two incipient species. (**B**) PCA and (**B’**) distance-based phylogenetic network of 22 male whole-genome SNPs from the admixed Maryland population. (**C-D**) *F*_*ST*_ values for *C. philodice* vs *C. eurytheme* plotted against recombination rate (**C**), and Manhattan plot (**D**). Red indicates windows with above-median recombination rate and in the 95^th^ percentile of *F*_*ST*_, including on the Z chromosome (asterisk). (**E**) QTL analysis of presence/absence of UV in 252 male offspring from F2 and Backcrosses. (**F**) Genotype plot for the whole Z chromosome with resequencing data from 23 individuals. Each row is an individual, and each column is a color-coded SNP. (**G**) Annotation of the *U* locus zero-recombinant interval (box) and surrounding region.

## Results

### The Z chromosomes define species barriers in secondary sympatry

To test a putative role of the Z chromosome as a barrier locus, we conducted a genome scan on 22 males from a sympatric population in Maryland, where *C. eurytheme* settled in 1927 (17). Two individuals were identified as probable recent hybrids and excluded from further analyses. We retained 11 UV, orange males and 9 non-UV, yellow males which formed two discrete clusters based on genome-wide SNP clustering by PCA (**Fig. 1C, Fig. S2, and Table S1**). The Z chromosome showed a large increase in genetic differentiation when compared to autosomes (**Figs. S3-4 and Table S2**), with a Z:A ratio of 12:1, the highest sex-chromosome to autosome ratio of *F*_*ST*_ reported from a whole genome dataset (7, 18). Heterogeneous landscapes of genomic differentiation can be explained by local barriers to gene flow or by linked selection in regions of low recombination (7, 19). To parse these two phenomena, we highlighted the top 5% windows of *F*_*ST*_ and an above-median recombination rate, thereby identifying three regions — two narrow autosomal *F*_*ST*_ peaks, and a 2.5 Mb portion of the Z chromosome (**Fig. 1C, D**). These data show that while only a restricted set of autosomal regions are likely under selection in each population, a large fraction of the Z chromosome is refractory to gene flow in a pattern consistent with a causal role in reproductive isolation. In other words, while Z chromosomes can recombine when hybrid F_1_ males are produced in the lab, this rarely persists in the field, presumably due to the aforementioned isolating mechanisms that select against hybrids.

In addition, nucleotide diversity (*π)* was depressed on the Z chromosome in both populations and divergence (*d*_XY_) was elevated, supporting the inference that the Z chromosome is highly differentiated (**Fig. S3C-D’**). In most scenarios one expects *π* on the Z chromosome to be 75% of *π* on the autosomes (20). For *C. eurytheme, π*_*Z*_*/π*_*A*_ was 0.751 ± 0.118, matching this expectation (**Fig. S3E**). However for *C. philodice, π*_*Z*_*/π*_*A*_ was 0.532 ± 0.105, meaning *π*_*Z*_ was lower than expected, which could reflect a recent selective sweep in *C. philodice*.

### The male mating signal polymorphism maps to *bab*

The large-Z effect results in extended non-recombining haplotypes in the natural population that prevent association mapping of trait variation (**Fig. S5**). To gain further resolution on the genetic basis of the polymorphic UV signal, we turned to linkage mapping from controlled hybrid crosses. F_2_ and backcross (BC) broods showed Mendelian, recessive segregation of the UV state among male offspring (**Fig. S6**). We genotyped 484 recombinant males and females using 2b-RAD sequencing, scored UV among the 252 genotyped males, and identified a LOD (logarithm of the odds) interval on the Z chromosome (**Fig. 1E and Fig. S6**). We resequenced individuals with recombination events around the *U* locus and refined a 352 kb zero-recombinant window with 18 annotated genes (**Fig. 1F, G, and Tables 4-5**). This mapping interval includes the 5’ intergenic region, promoter, and first exon of the gene *bric a brac* (*bab*), a salient candidate gene as it encodes a transcriptional repressor of male-limited traits in *Drosophila* such as abdominal pigmentation and sex combs (21–27).

### Bab expression marks non-UV scale cells

To test a role in the regulation of *Colias* male UV, we characterised the expression and developmental functions of *bab* during color scale formation. Butterfly wing scales are macrochaete derivatives that each protrude from a single epidermal cell during pupal development (28). UV scales found in the subfamily Coliadinae are a derived scale type characterised by dense longitudinal ridges; each scale forms a multilayer of chitinous lamellae that selectively reflects UV light by the coherent scattering of incident light (29–32). UV scales are specific to the dorsal wing surface of *C. eurytheme* males, and cover the top of non-UV ground scales (**Fig. 2A, B**).

**Fig. 2.**
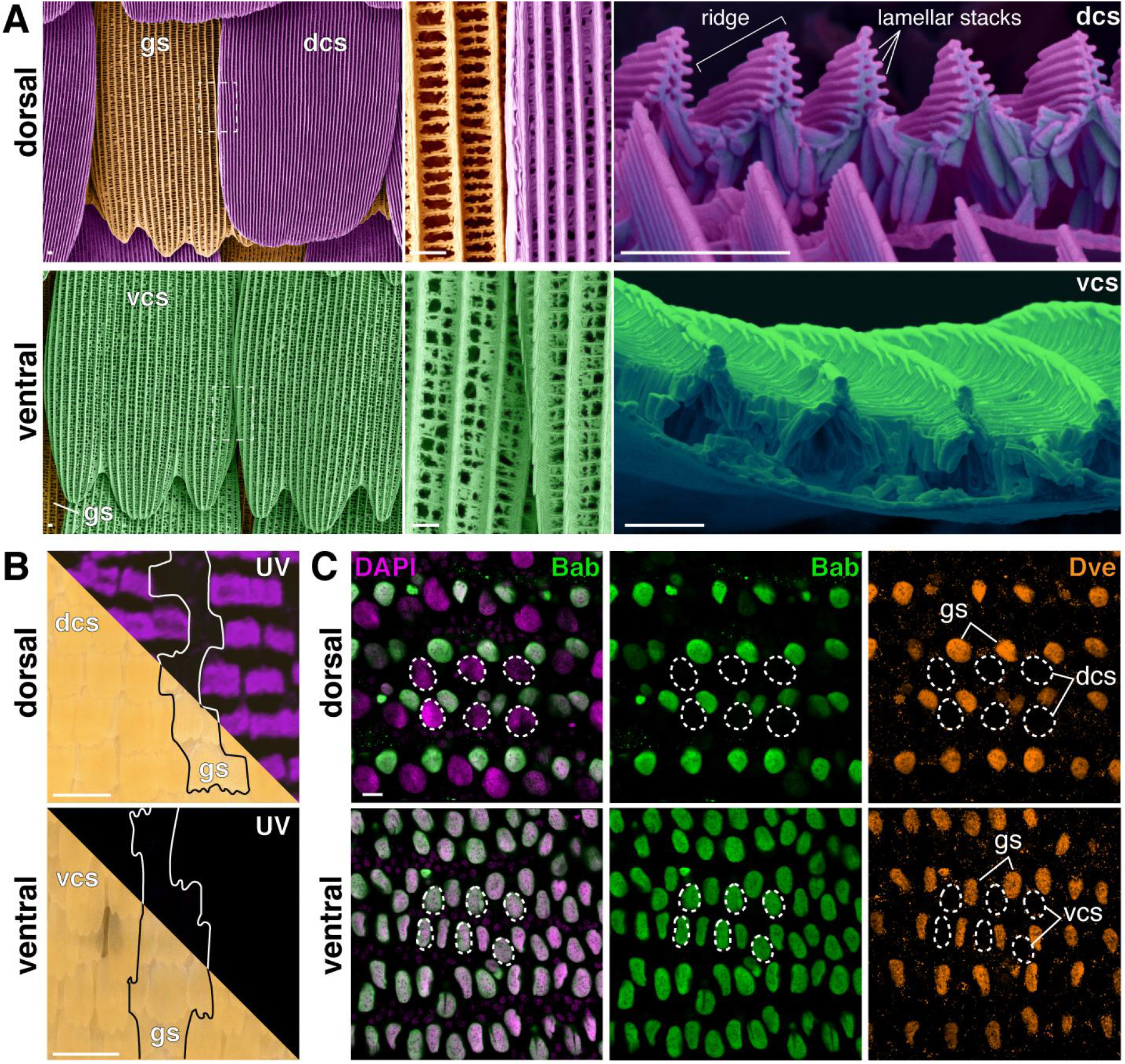
Bab negatively correlates with UV scale precursors in *C. eurytheme* male wings. (**A**) Pseudo-colored SEMs highlighting the ultrastructural differentiation of the UV+ dorsal cover scales (dcs, magenta), relative to UV-ground scales (gs, orange), and UV-ventral cover scales (vcs, green). (**B**) Microphotographs of adult *C. eurytheme* male wing surfaces in the visible and UV ranges. Line: damaged areas exposing UV-ground scales. (**C**) Immunofluorescent detection of Bab (green) in all UV-precursors at 46% pupal development. Magenta: DAPI (nuclei) ; orange: Dve ; circles: cover scale nuclei. Scale bars: A = 2 μm ; B = 100 μm ; C = 10 μm.

Immunofluorescence at the onset of scale emergence reveals Bab expression in the nuclei of all non-UV scale cell precursors (**Fig. 2C, Fig. S7, and Movie S1**) regardless of the scale layer (ground, cover), pigment fate (yellow pterins, black melanin), wing surface, and sex. Remarkably, Bab is expressed in ground scales and is absent in the UV cover scales on the dorsal male wing surface. Thus, Bab is negatively correlated with the UV scale type in *C. eurytheme*: it is consistently expressed in all scale cells fated as non-UV except in the wing cover scales of *C. eurytheme* males. Where it is not expressed, these scales develop layered nanostructures specifically capable of UV-iridescent reflectance. This inactivation in the sexually dichromatic pattern suggests a repressor function, analogous to the expression of Bab in the *Drosophila* abdominal epithelium (21, 23, 27).

### CRISPR knock-outs yield ectopic UV-iridescence

To directly test this model, we generated CRISPR-mediated loss-of-function mutations targeting the first exon of the *bab* coding sequence. We collected *C. eurytheme* and *C. philodice* females and microinjected eggs within 7 hrs post-fertilization. G_0_ *bab* crispants showed mosaic phenotypes of high penetrance (51 out of 63 surviving adults), with a widespread gain of UV in both males and females of both species, including ventral surfaces (**Fig. 3 and Figs. S8-13**). Both pterin and melanin pigment scales, and both cover and ground scales differentiated into UV scales following *bab* knockout (KO). Female-specific effects on pigmentation were also noted **(Fig. S11**). These loss-of-function assays show that Bab represses the UV identity in all non-UV scale precursors regardless of wing surface, sex, or species. Male UV iridescence is widespread in the *Colias* genus (32), suggesting that the *C. philodice* absence of UV is due to a secondary loss of Bab repression. The reappearance of UV in *C. philodice bab* crispants (an atavism) implies that the underlying network for producing UV scales is still present in this species.

**Fig. 3.**
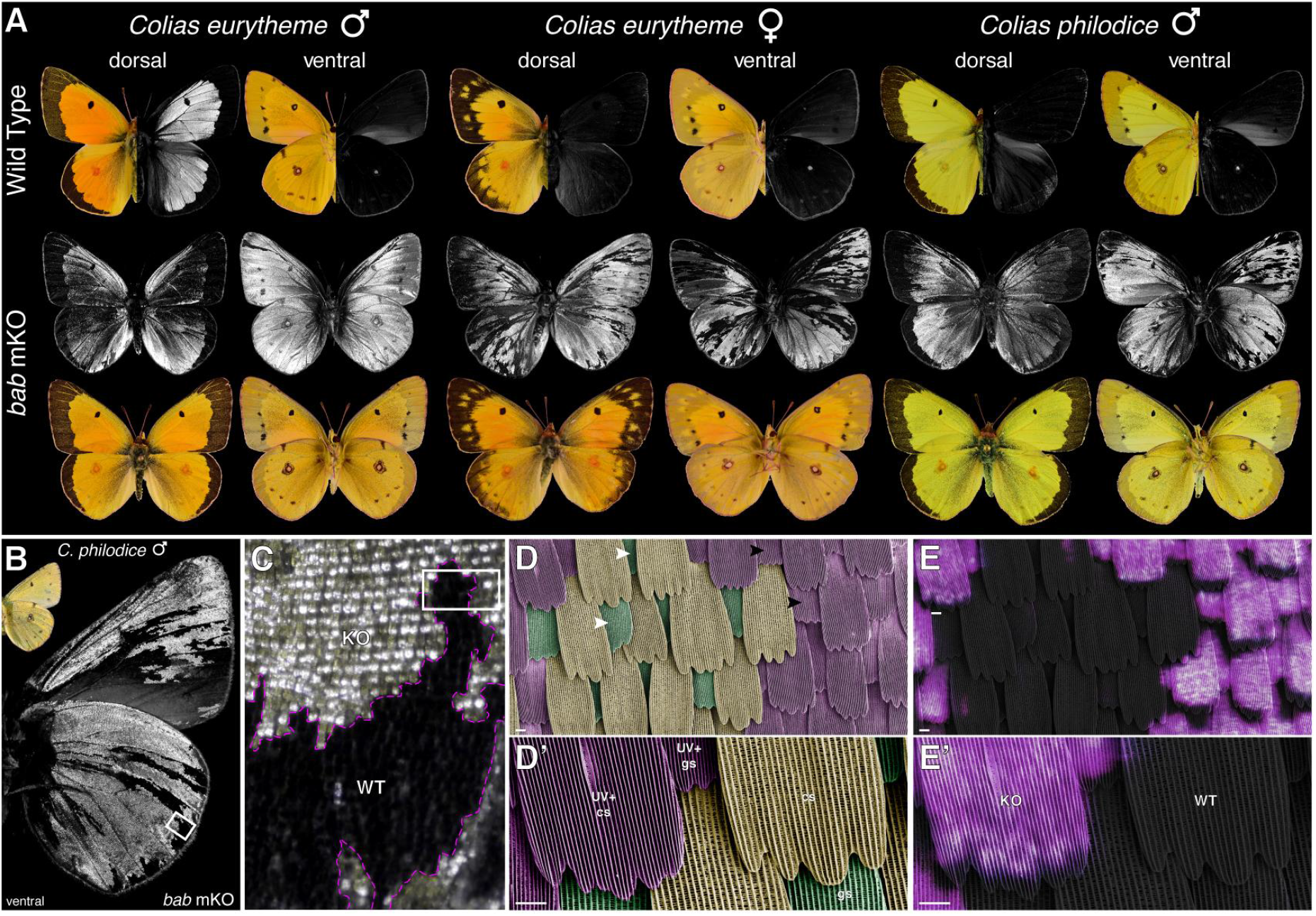
Bab represses UV scale fate. **A**. G_0_ phenotypes resulting from CRISPR mosaic KOs (mKO) targeting the first exon of *bab*. Gain of UV-iridescence was observed across both species and both sexes, including ventral wing surfaces. **B-C**. Magnified view of male *C. philodice* wings featuring extensive ectopic iridescence following *bab* mKO. **D-D’**. False-colored SEM view of a mosaic region (box in C), with complete transformation of wild-type cover (cs, orange) and ground scales (gs, green) into UV-iridescent scales with dense longitudinal ridges. Arrowheads: examples of ground scales in a wild-type state (white) and in a *bab* mutant clone. **E-E’**. Correlative light and electron microscopy featuring a superimposed view of UV-reflected light (magenta) on an SEM view. Both ground and cover scales from *bab* mutant clones are UV-iridescent. Scale bars = 10 μm.

## Discussion

### Bab is a repressor of male dimorphic traits

Our combined linkage mapping, expression and functional assays of UV iridescence show that *1)* allelic variation of *bab* causes a male-specific mating signal difference between two incipient species; *2)* Bab expression is sexually dimorphic in *C. eurytheme*; and *3)* Bab functions as a repressor of the dimorphic trait. This repression of a male-specific feature is analogous to the expression and function of Bab paralogs in *Drosophila* sex-comb formation, abdominal pigmentation, and gonadal stem cell niches (21–23, 26, 27, 33, 34), with a male-specific repression in the presumptive cells forming the male feature, and loss-of-function resulting in gain or expansion of the male state. Bab is thus a major player in the development of sexually dimorphic features, and might have an ancestral function in the repression of male-specific development at the root of butterflies and flies. It will be critical to study the versatility of its functions comparatively in order to better understand how sexual forms evolve. As knockdowns and knockouts are increasingly amenable in new organisms, testing the repressive nature of Bab should be feasible and may yield a wide range of trait gain or masculinization phenotypes. Further work could also explore how the dimorphic (*C. eurytheme*) vs. monomorphic (*C. philodice*) expression of Bab is achieved. A potential regulator could be the transcription factor Doublesex, an integrator and master selector gene of somatic sexual identity in insects, including butterflies (35–38). In fruitflies, the DsxM male isoform of Doublesex directly represses Bab via a *cis-*regulatory element in the first intron of *bab1*, thus derepressing abdominal pigmentation, while the DsxF female isoform activates Bab, repressing pigmentation (24, 28). Extrapolating from this precedent, it will be interesting to test if DsxM similarly represses Bab in *C. eurytheme* cover scales, and if its binding sites have mutated in *C. philodice*.

### *bab* is a genetic hotspot of sexual phenotypic variation

There is replicated evidence that *cis*-regulatory evolution of *bab* directly causes sexual trait divergence in flies and Lepidoptera (butterflies and moths), making it a genetic hotspot of phenotypic variation (39). In *Drosophila*, linkage mapping studies have shown that regulatory alleles of the *bab* locus (*bab1/2* recent paralogs) explain natural variation in dimorphic pigmentation and ovariole number (23, 26, 40). In corn borer moths (*Ostrinia nubilalis*), male response to a polymorphic female pheromone blend is driven by a Z-linked regulatory variation in the first intron of *bab* (41). This same intron is within the *Colias U* locus interval. Thus, the 5’ portion of the lepidopteran *bab* locus likely underlies variation in a male olfactory preference in recently diverged *Ostrinia* species, as well as in a male visual signal in *Colias*. This leads us to propose that the *bab* locus is a hotspot for the evolution of reproductive isolation, driving species divergence and maintenance across Lepidoptera. Factors related to the function, regulation, and genomic location of *bab* may have predisposed it to the tuning of sexually selected traits. First, repressors have drastic effects on cis-regulatory regions and efficiently suppress transcription (42, 43). In this sense, evolving new Bab binding sites in DNA enhancers might be a path of least resistance for optimizing gene expression subtractively, rather than by adding activator binding sites for other selector genes. There is already precedent for the *Drosophila* Bab integrating into a pre-existing network and sculpting sexual dichromatism (27, 44) consistent with this idea. Second, the published expression patterns of Bab show it integrates both spatial and sex-determination inputs (21, 24, 25, 34). Due to a hub-like position in gene regulatory networks, Bab is potentially an input-output gene that facilitates tissue-specific change (45, 46). The large 5′ intergenic and intronic regions of *bab* in *Colias* and *Ostrinia*, and of its sub-functionalized duplicates in *Drosophila*, together suggest a complex cis-regulatory landscape bearing a multitude of enhancers or silencers (47–49), enabling the evolution of precise, tissue-specific changes (50). Last, the location of Bab on the lepidopteran Z chromosome is relevant to the divergence of premating traits under the assumption that sex-chromosomes are more prone to the generation of reproductive isolation than autosomes (20, 51, 52). These possible generative biases will require further investigation among diverged *Drosophila* species, as well as in the *Colias* UV display and *Ostrinia* mate detection systems. Noneless, we speculate that the peculiar molecular function, regulation, and genomic location of Bab may collectively explain its propensity to fine-tune variation in sexual traits.

### Extreme Large-Z effect in an anthropogenic contact zone

Our genomic scan of differentiation focused on two incipient species that recently reunited due to human activity (alfalfa agriculture). We caught a remarkable signature of a large-Z effect (analogous to large-X) with widespread admixture across autosomes, and strong differentiation of the entire Z chromosome. This heterogeneous landscape of differentiation is the most pronounced identified so far from genome-wide data (18), supporting a role for sex-chromosomes as key drivers of reproductive isolation (20, 51). Collectively with the evidence that both premating and postmating isolating mechanisms are sex-linked in this system (12, 14, 15), these data highlight that species status in this system is determined by Z chromosomes, while autosomes are exchangable. While linked selection (*syn*. divergence hitchhiking) in regions of low recombination can sometimes explain such extensive blocks of divergence (7, 19), our linkage map did not indicate that this is the case in the *Colias* sympatric pair. The *U* locus in particular overlapped with a large region with above-median recombination in between-species crosses, but high *F*_*ST*_ indicates it is refractory to gene flow in wild populations, strongly implying that it acts as a prezygotic genomic barrier to gene flow rather than as a block of divergence hitchhiking. Overall, these data establish the anthropogenic contact zones of *Colias* butterflies as a promising system for the study of large-Z(X) effects in speciation with gene flow.

### Coupling of reproductive barriers

The data in hand suggests the *U* locus variation causes premating isolation by accurately displaying sex chromosome compatibility of courting males to *C. eurytheme* females. In summary, the *Colias* UV mating signal is polymorphic and recessive in areas of secondary contact, with both heterospecific and hybrid males lacking UV because they carry the dominant *U* allele of *C. philodice*. This allele drives uniform expression of Bab, preventing UV scale development with a dominant effect. Conversely, the recessive *C. eurytheme u* allele represses expression of Bab in dorsal male cover scales leading to UV fated scales. This Z-linked UV signal allows *C. eurytheme* females to choose Z-compatible mates, thereby prezygotically selecting against Z-linked hybrid sterility (3, 9, 10, 12–14). The recessivity of the trait allows the rejection of Z-heterozygous males and prevents a 25% cost in reproductive fitness (**Fig. S1**), an effect that would not be possible if Bab were an activator of UV, which would likely be dominant. In this way, a large-Z effect that couples pre- and postmating barriers likely drives reproductive isolation in these butterflies (5, 18), akin to the genetic architectures of speciation in other species (16, 53), and previously conceptualized as an “indicator” mechanism that enables assortative mating (3). Further work is needed to decipher other putative barrier loci on this chromosome. In any case, these findings highlight how a simple genetic switch for a mating cue can influence the origin of species and the maintenance of biodiversity.

## Experimental Procedures (also see Supplementary Text)

### Genomic scans of differentiation in sympatric males

Samples originated from a large syntopic population at an alfalfa farm in Buckeystown, MD. Whole-genomes from 24 males were resequenced at 14.3x mean coverage, aligned to the *C. eurytheme* reference genome assembly (54), and used in a population scan analysis pipeline (55).

### Linkage map

Interspecific crosses consisting of an F2 and two back-cross broods generated 528 recombinant individuals of known pedigree, sex, and UV phenotype. 2b-RAD sequencing was used to genotype 484 males and females in a HiSeq 4000 SE50 run, using the *Bcg*I enzyme and adapters yielding a 16-fold representation reduction (56). Genotypes were used to build a linkage map as described elsewhere (54). Select individuals were resequenced at a 15x mean coverage to narrow the *U* locus interval.

### Bab expression and loss-of-function assays

A custom rabbit polyclonal antibody was generated against the N-terminal 1-365 residues of the *C. eurytheme* Bab protein, and used with a guinea pig anti-Dve (57) for whole-mount immunofluorescence in pupal wings. Heteroduplex mixes of Cas9-NLS and two equimolar sgRNAs (500:125:125 ng/μL) were injected in 1-7 hrs syncytial embryos for targeted mutagenesis of the first exon of *bab*.

### CRISPR-mediated KO of *bab*

Two overlapping guide RNAs were designed targeting the first exon of bab within the U-locus. Heteroduplex mixes of Cas9:sgRNA1:sgRNA2 (500:125:125 ng/μL) were prepared and microinjected in butterfly syncytial embryos 1-7hrs AEL. Eggs incubated for two days at 28°C then placed on vetch sprouts aged 7-14 days at an average greenhouse temperature of 24°C. Two rounds of injections were successfully performed under these conditions resulting in 51 of 63 adults displaying crispant phenotypes.

### UV photography

UV-photography was performed using a full-spectrum converted Lumix G3 camera, mounted with Baader U-Venus filters and UV-transmitting lenses, under the illumination of blacklight bulbs or 365nm LEDs.

## Data and Code Availability

Whole-genome sequencing data are available in the Sequence Read Archive (www.ncbi.nlm.nih.gov/sra) under BioProjects PRJNA663300, PRJNA719421, and PRJNA723900. SNP calling, genotyping data, and computer code are available from the Dryad digital repository (55).

## Acknowledgements

We thank B. Wang, S. Barao, W. Watt, and R. Canalichio for butterfly rearing, access and expertise, R. Rogers, S. Van Belleghem and P. Rastas for bioinformatics assistance, C. Brantner and C. Day for SEM, M. Matz for 2b-RAD protocols, L. Livragh for vector graphics, and M. Perry for the Dve antibody. We also thank core facilities at the GWU, UT Austin, and U. Maryland-Baltimore for generating sequence data, and the GWU HPC team for providing computing infrastructure (58). This work was supported by the NSF awards IOS-2108227 and IOS-1656553, and the Swedish Research Council award 2017-04386.

## Supplementary Text

### Genomic scans of differentiation in sympatric males

#### Genome resequencing

We collected and resequenced 24 males (**Table S1**) from a syntopic population found in organic alfalfa fields (Hedgeapple Farm) in Buckeystown, MD. DNA was extracted from thoracic tissues using the Qiagen DNeasy Blood & Tissue Kit, RNAse-treated, and used to prepare a multiplexed sequencing library with the TruSeq PCR-free DNA protocol. The resulting library was sequenced in a first test run using an Illumina NextSeq500 Pair-End 2×75 bp run, and then reloaded in a second Illumina NextSeq500 Pair-End 2×150 bp run, yielding a total of 14.3x coverage per individual on average. Sequencing reads are accessible in the NCBI SRA under the project accession number PRJNA663300.

#### Alignment, genotyping, and genome-wide analyses

Reads were aligned to the *C. eurytheme* reference genome, derived from a California individual (54). Samples were aligned with BWA-MEM using default parameters (Li, 2013), and variants called using GATK v4.1 with tools HaplotypeCaller and GenotypeGVCFs using default parameters (59). Variant sites were accepted if they were biallelic and the quality (QUAL) value was ≥30. SNPs were phased with Beagle 4.1 (60). Phased SNP variants were used to perform principal component analysis (PCA) using the Eigensoft module SmartPCA (61), and dendrograms were computed using code by Simon H Martin (62) and visualised with SplitsTree (63).

As *C. eurytheme* and *C. philodice* are known to hybridize when found in sympatry at high densities, we sought to determine if any of our sequenced samples were recent hybrids. The PCA showed two main clusters, one containing all *C. eurytheme* and the other containing most *C. philodice*, with clear separation between species along PC1 **Fig. S2A**). Two samples (numbered *03*, phenotypically identified as *C. philodice*, and *04*, phenotypically identified as *C. eurytheme*), formed a separate cluster. Distance matrix dendrograms grouped these two individuals with the *C. eurytheme* clade, with each other as the closest relative. We determined that these individuals are likely recent hybrids and therefore excluded them from genomic analyses. We re-analysed the data with the likely hybrids removed, and retrieved two clusters with strong separation along PC1. When looking at all genomic data, two *C. eurytheme* individuals were separated on PC2, but were nested within the rest of the *C. eurytheme* individuals on PC1 and in the distance matrix dendrograms (**Fig. S2B**). Likewise, when computing a PCA for just the Z chromosome, two clusters formed with clear separation on PC1, but one individual (numbered 26) was separated by PC2 (**Fig. S2C**). For the rest of the analyses herein, the two sample clusters were used to define each population.

#### Genome scan using population genetic statistics

We computed population genetic statistics in windows of 100 kb with a sliding overlap of 10 kb (55, 62). Mean genome-wide divergence indicated by fixation index (*F*_*ST*_) between species is low, with peaks of high divergence on autosomes (**Fig. S3A, B**). The Z chromosome shows considerably higher mean divergence than autosomes, consistent reduced gene flow of the sex chromosome caused by hybrid inviability, and is also consistent with the observation that wing patterning traits in *Colias* are clustered on the Z chromosome (13). The elevated Z/A divergence ratios we report for *Colias* are best exemplified when compared to lower Z/A values reported for *Heliconius* species, that were calculated using similar sliding window methods and sample sizes as in our analysis (64).

#### Recombination rate in interspecific crosses

Genome-wide recombination rate was computed from the linkage map using the R package MareyMap (65) using a loess smooth with a span of 1 Mb and a degree of 1. A cM/Mb value was extracted for 100 kb intervals along the entire genome (**Fig. S3F**).

### Linkage mapping of UV variation in males

#### Mapping crosses

Interspecific crosses consisting of an F2 and two back-cross broods (named broods 75, 49, and 79 respectively) were reared in the laboratory of Adam Porter in 2000 and described in a previous publication (13). Briefly, the offspring of wild-caught females from the vicinity of Amherst and Sunderland, MA were reared in the laboratory for species assignment, and used for controlled crosses over two generations where all brood pedigrees and phenotypes were recorded. UV-iridescence showed binary presence-absence and Mendelian segregation ratios that confirmed the recessive and sex-linked inheritance the *U* locus (**Fig. S5**). Crossing-over is absent from lepidopteran females but recombination is expected in the hybrid F1 fathers (ZZ) of those three crosses, which are thus informative for the mapping of the male-expressed and Z-linked *U* locus. A total of 528 adult F2 and BC butterflies were frozen at -80°C after emergence until further analysis.

#### DNA extraction

Frozen adult specimens from the mapping broods were each given an identifier, and sequentially processed for genotyping and phenotyping. Wings of each individual were removed and dispatched into glassine envelopes, the dorsal half of the thorax was excised and transferred to the well of a 96 Collection Microtube Rack (Qiagen, USA) with a 3.5mm stainless steel bead, and the rest of the body was stored in an individual tube for frozen storage. To prevent cross-contamination of DNA samples, each individual was processed on a new aluminium foil surface and tools were cleaned in 5% bleach, rinsed in distilled H_2_O, and dried with EtOH 70% between samples. Thoracic tissue samples were mixed with 400 μl of BeadBashing Buffer (Zymo Research, USA), and homogenised in the collection racks on a MM400 mixer mill (Retsch, Germany) fitted with a TissueLyser 2 × 96 Adapter Set (Qiagen, USA) for 4 min at 30 Hz. Following centrifugation, the supernatant from those homogenates was DNA-extracted while maintaining a 96-well microplate format using the Quick-DNA 96 kit (Zymo Research, USA), eluted in 50 μL of DNA Elution Buffer, and quantified with a Qubit dsDNA High-Sensitivity kit.

#### Preparation of the 2b-RAD sequencing library

The mapping broods resulted in 497 DNA samples spread across six 96-well plates (55). We used 2b-RAD genotyping with a multiplexed strategy to pool all the samples into a single sequencing library (56, 66). 2b-RAD sequencing uses Type IIb restriction enzymes that cleave DNA on both sides of their targets. We used the *Bcg*I enzyme to generate 34 bp fragments flanked by barcoded sequencing adapters (**Table S3**). Adapters terminated on each side end with NG-3’ instead of NN-3’, resulting in a 16-fold reduction in the representation of all restriction sites. Samples were dispatched in new microplates with 200 ng of DNA per sample (25 μL, normalised at 8 ng/μL), digested with *Bcg*I, heat inactivated, and purified with the ZR-96 Oligo Clean & Concentrator kit (Zymo Research, USA) in a final elution volume of 12 μL. The purified restricted DNA was then ligated with the generic 5’ adapter (*5ILL-NG*) and with a different 3’ adapter (*3illBC1-12*) in each column, before heat inactivation and pooling of the 12 plate columns into a single strip of 8 rows. This product was then PCR amplified for 14 cycles with standard Illumina adapters (IC1-P5, IC2-P7, and NEB Universal PCR primer) as well as one of 48 unique barcoded adapters (*NEBNext Multiplex Oligos Index Primers Sets 1-3*), yielding a visible band of about 170 bp on an agarose gel. Those products were quantified by fluorometry and pooled into a single library. A sequencing facility purified the target product of 170 bp using a BluePippin instrument equipped with 3% Agarose DNA cassettes (Sage Science, USA), quality checked, mixed with a 20% *PhiX* phage library, and sequenced using a Illumina HiSeq 4000 SE50 run.

#### Linkage mapping of the U locus

The linkage map was built in LepMap3 as previously described (54). Data were imported to R/qtl as a four-way cross (55, 67). First, we performed a QTL analysis with the male samples, treating the presence or absence of UV as a binary trait and performing 1000 permutations to determine a confidence threshold. A strong QTL peak was identified on the Z chromosome, with an extended LOD interval. As the trait is Mendelian, we then looked for regions on the Z chromosome that showed an inheritance pattern that was completely consistent with phenotype (zero recombinant window), and identified a number of individuals with recombinations around marker 41 (**Fig. S6**).

#### Resequencing of recombinant individuals

To identify the precise breakpoints in these recombinant individuals, we resequenced their undigested genomes at 15x mean coverage using an Illumina NovaSeq S1 2×150bp run, along with three brood grandmothers and an F1 male (**Table S4**). Raw reads were deposited in the SRA under the BioProject PRJNA719421. These samples were aligned, variant-called and phased as above. We filtered Z Chromosome SNPs for sites where both the *C. philodice* and *C. eurytheme* grandmothers had alternate alleles and the F1 male was heterozygote. The data for the Z chromosome was visualised using *genotype_plot* (55, 68, 69), and breakpoints manually identified (**Fig. S4**). This visualisation includes the individuals from Buckeystown, MD; note that individual 26, which in the Z chromosome PCA was separated on PC2 from the other *C. eurytheme* (**Fig. S2C**), appears to contain a large heterozygous block of *C. philodice* haplotype on the 3’ end of the Z chromosome. We refined the *U* locus (**Fig. 1E**) to the interval between the breakpoints in individuals *00-79-099* and *00-49-013*, which runs between positions *Sc000009*: 5173306-5525979. There are 18 annotated transcripts within or overlapping this interval in the *C. eurytheme* GFF (**Table S5**) including the first exon and part of the first intron of the gene *bab*.

#### Protein coding alignment of Bab

Variants in the protein coding sequence of *bab* were extracted for each resequenced individual (**Tables S1, 4**), converted to FASTA format using *bcftools*, and aligned with *MAFFT* in Geneious. No amino acid or nucleotide variants were fixed between UV-iridescent and non-iridescent males, though some polymorphism exists within these two populations. Nucleotide and amino acid alignments are available on DRYAD (55). There are no coding variants in the first exon, the only coding exon of *bab* situated within the boundaries of the *U* locus, implying that the *bab* variation responsible for the UV interspecific polymorphism is of *cis-*regulatory nature.

### Expression and loss-of-function assays

#### Antibodies

The production of the custom rabbit anti-Bab polyclonal antibody was outsourced to a manufacturer (GenScript Biotech, NJ). A sequence encoding His-tagged amino-acid residues 1-365 from the *C. eurytheme* annotated protein was cloned in the *pET-30a* plasmid, and purified from bacteria with a His-tag before immunisation of two rabbits, from which polyclonal sera were affinity purified before use. The guinea pig anti-Dve antibody was a kind gift from Michael Perry (UC San Diego), and recognises a butterfly homolog of Defective proventriculus (57). Secondary antibodies used include conjugated AlexaFluor647 anti-Rabbit IgG (Life Technologies, CA) at 1:100 dilution, and conjugated AlexaFluor555 goat anti-guinea pig IgG (Abcam, UK) at 1:400 dilution.

#### Immunostainings

*Colias spp*. larvae were reared on 1-3 week old sprouts of Lana woolypod vetch (*Vicia villosa*) grown on hemp mats, and monitored for pupation throughout the day. Pupal developmental stages were temperature-dependent, with average adult emergence of 192 h APF (hours after pupa formation) at 24 °C and 144 h APF at 28 °C. Pupae were dissected at stages spanning 30-45% developmental time in phosphate buffer saline (PBS), fixed for 12 m at room temperature in formaldehyde 4% (PBS, EGTA 2mM), washed in PBS 0.1% Triton X-100 (PT 0.1%), blocked with PT 0.1% with 0.5% bovine serum albumin for a minimum of 30 m, incubated with primary antibodies overnight at 4 °C (anti-Bab, 1:100 dilution ; anti-Dve, 1:400 dilution), washed, incubated with secondary antibodies at room temperature for 2 h, washed, incubated in 50% glycerol, PBS, 1 μg/ml DAPI for nuclear counterstaining, mounted on glass slides with 70% glycerol under a #1.5 thickness coverslip, and sealed with nail varnish before imaging.

#### Confocal imaging

Stacked acquisitions were made using an Olympus FV1200 inverted laser scanning confocal microscope mounted with a 60x Apochromat oil-immersion objective (PLANAPO, 1.42NA), and equipped with a laser line for excitation at 405 nm (DAPI), 488 nm (Phalloidin-OregonGreen), 555 nm (anti-Bab / AlexaFluor555) and 647 nm (Dve / AlexaFluor647). Z-stacks were visualised in 2D using FIJI/ImageJ and in 3D with Imaris 3D/4D visualization software (Bitplane AG, Switzerland).

#### CRISPR-Cas9 mediated knockouts

Two unique sgRNA targets were designed in the first exon of *bab* in a region void of SNPs (*Ceu_bab_sgRNA1* target: 5’-ACTGTTGGGGCGAGCCGGG-3’ ; *Ceu_bab_sgRNA2* target: 5’-CGGCGGGCCCGGCTCCTCG-3’). Heteroduplex mixes of Cas9:sgRNA1:sgRNA2 (500:125:125 ng/μL) were prepared and microinjected in butterfly syncytial embryos as previously described (70). Continuous captive rearing of *C. eurytheme* and *C. philodice* is challenged by the sensitivity of the larvae to viral disease (71), and we thus used wild-caught females (Hedgeapple Farm, Buckeystown, MD) to obtain eggs from both species. Females were placed in 28 cm foldable cages with a high density of fresh alfalfa cuts in water, and under F54T5HO fluorescent tubes. Eggs were surface sterilised in 5% benzalkonium chloride, rinsed, dried, positioned on double-tape, microinjected 1-7 hrs after egg laying, placed in a plastic tupperware with a moist paper towel at constant 28 °C for 24-48 hrs and placed on vetch sprouts aged 7-14 days at an average greenhouse temperature of 24 °C. Trays of 8-21 days old vetch were added as necessary and surviving individuals reached adulthoods in an average of 30 days after egg laying. Larval mortality was high due to viral disease including in uninjected batches. Two rounds of *bab* CRISPR injections were performed : the first round resulted in 50 emerged G_0_ adults (997 eggs injected) who all showed mosaic, ectopic gains of UV-iridescence ; the second round resulted in 13 G_0_ adults (136 eggs injected) of which 11 showed positive results and 2 appeared wild-type.

### Phenotyping and imaging

#### Photography in the ultraviolet and visible ranges

We combined digitisation in the visible color range and UV-photography to phenotype the wings of the individuals used in this study. For UV imaging, wings and specimens were illuminated by CFL BlackLight 13 Watt T3 Spiral Light Bulbs (General Electric, USA), and photographed using a modified Panasonic/Lumix G3 with full-spectrum UV/VIS/IR conversion purchased from an online retailer (Infraready, UK). For selective imaging of the UV reflectance, this camera was mounted with a custom-modified 75mm UV-transmitting lens (source : eBay user Igoriginal), stopped at f/11, on an Adjustable T-to-M4/3 Mount Lens Adapter, and fitted with a Baader 2’’ U-Venus-Filter (Baader Planetarium, Germany), which transmits 80% of light above 350 nm but blocks all visible light in the visible range above 400 nm. For UV-microphotography of scales, the same camera was fitted with a Baader 1¼’’ U-Venus-Filter, and mounted with UV-transmitting lenses that allowed various levels of magnification, including an EL-NIKKOR 50mm f/2.8N lens reverse-mounted on extension bellows, an Olympus UPLANFL N 10x/0.30NA objective, and a Nikon CFI S Plan Fluor ELWD 20X/0.45NA objective. At the highest magnification ranges, best results were obtained with increased illumination using an U301 365nm Nichia UV LED Flashlight (MTE, China). Pinned specimens were photographed in the visible range using a Nikon D5300 camera mounted with a Micro-Nikkor 105mm f/2.8G lens on a StackShot rail, and focused-stacked using Helicon Remote and Helicon Focus (Helicon Soft, Ukraine). A Keyence VHX-5000 digital microscope was used to generate stitched high-resolution images of wing patterns using a VH-Z00T lens at 50X magnification and a VH-Z100T lens at the 300X magnification. The excised wings of all sequenced specimens were scanned on an Epson Perfection V600 scanner at 6,400 dpi resolution next to a color reference card.

#### Scanning electron microscopy

For surface imaging of ground and cover scales, regions of interest were excised and mounted on SEM stubs with double-sided carbon tape and sputter coated with 10 nm of gold. Images were acquired on a FEI Teneo LV FEG SEM, using secondary electrons (SE) and an Everhart-Thornley detector (ETD) with a beam energy of 2.00 kV, beam current of 25 pA, and a 5 μs dwell time. Individual images were stitched using the Maps 3.19 software (ThermoFisher Scientific). To document scale ultrastructure, regions of interest were excised and cryo-fractured following previous recommendations (72–74). Excised wing sections were placed target-side down onto a silicon wafer and secured with foam board, glassine, and a binder clip before submersion in liquid nitrogen, and immediately cut with a fresh ceramic-coated microtome blade. After drying, individual cut scales were placed on copper tape using an eyelash tool, such that the cut edges were approximately parallel to and overhanging the tape edge. The copper tape was bent to 90° and placed on a stub so that the scales’ cut edges faced upwards, *i*.*e*. normal to the stub surface, and secured with additional copper tape. Stubs were sputter coated with a 10 nm layer of gold, and imaged at 5.00 kV / 6.3 pA and a 10 μs dwell time.

#### Correlative light and electron microscopy

Wing samples were cut and mounted on an SEM and UV-imaged with the reverse-mounted El-Nikkor 50mm f/2.8N lens, allowing an initial capture of UV-iridescence across the sample. The region of interest was then imaged at the 700X magnification setting on the Keyence VHX-5000/VH-Z100T microscope for registering dimensions in the visible range, and imaged in the UV range, with best results using the Nikon CFI S Plan Fluor ELWD 20X objective, acquisitions of 20 images with increments of 3 μm using the Stackshot Rail, and depth-map stacking (Method B) in Helicon Focus. Stubs were then sputter-coated and stitch-imaged at 3,500X, with the navigation camera recording the precise position of the acquisitions. SEM, visible, and UV images were aligned in Adobe Photoshop, and in final renderings UV images were set-up with an opacity around 70% superimposed on SEM using the Hard Light blending mode.

## Supplementary Figures

**Fig. S1.**
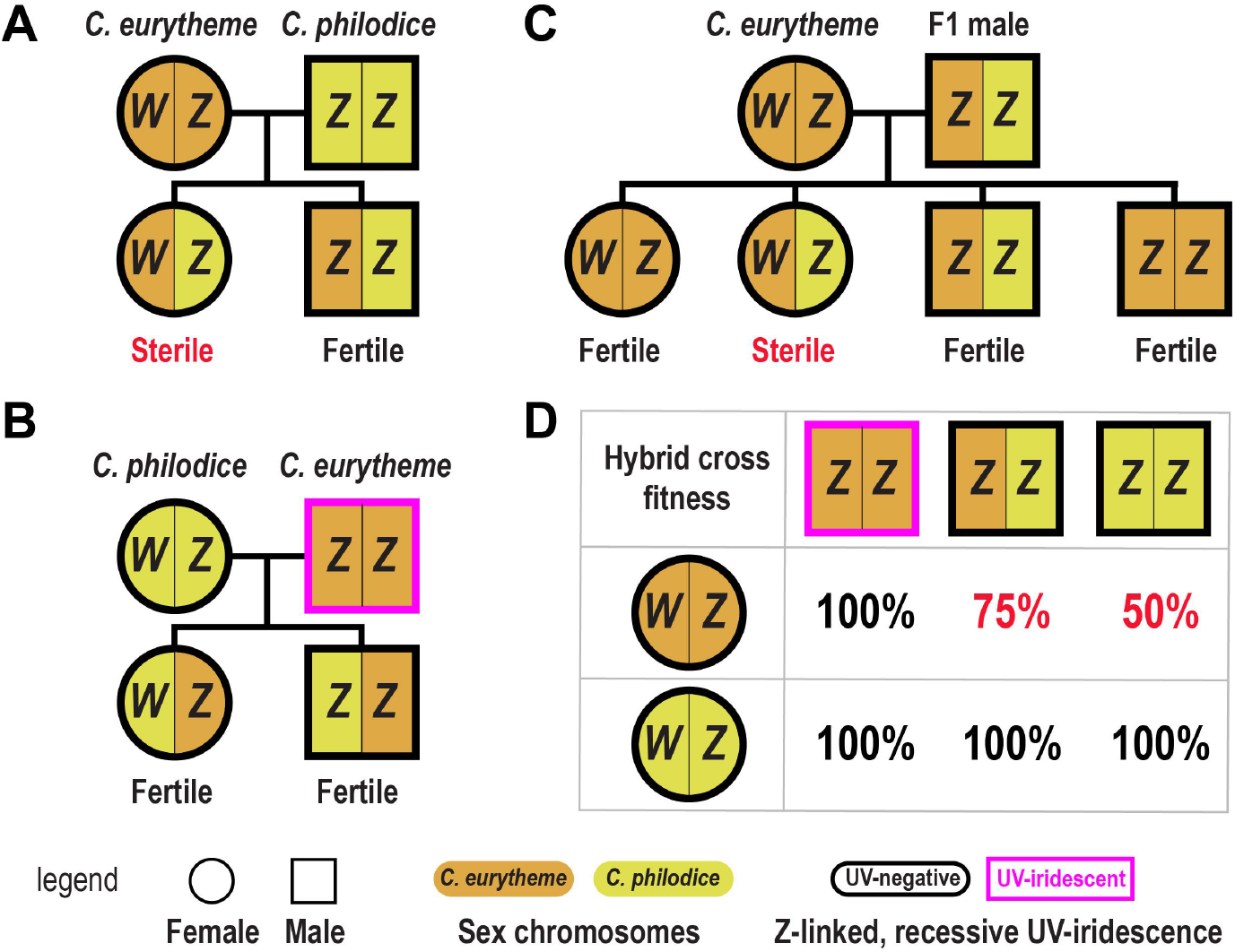
Female hybrid sterility in *C. eurytheme x philodice* butterflies. **A-C**. Data summary from Taylor and Grula (12) illustrating the pattern of Z-linked female hybrid sterility among sympatric sulphur butterflies. **D**. Parental crosses involving a *C. eurytheme* W mother and *C. philodice* Z sperms suffer a reproductive fitness reduction (red), but these incompatibilities can be avoided by assortative mating of *C. eurytheme* females with UV-iridescent males homozygous for the *C. eurytheme* Z chromosome. Sterile females develop normally but undergo ovarian failure, a phenomenon that could arise from incompatibility of the *C. eurytheme* matrilineal elements (W chromosome, mitochondrial genome) with the *C. philodice* Z chromosome during oogenesis, or from a conflict between the *C. philodice* Z and an autosomal *C. eurytheme* incompatibility locus.

**Fig. S2.**
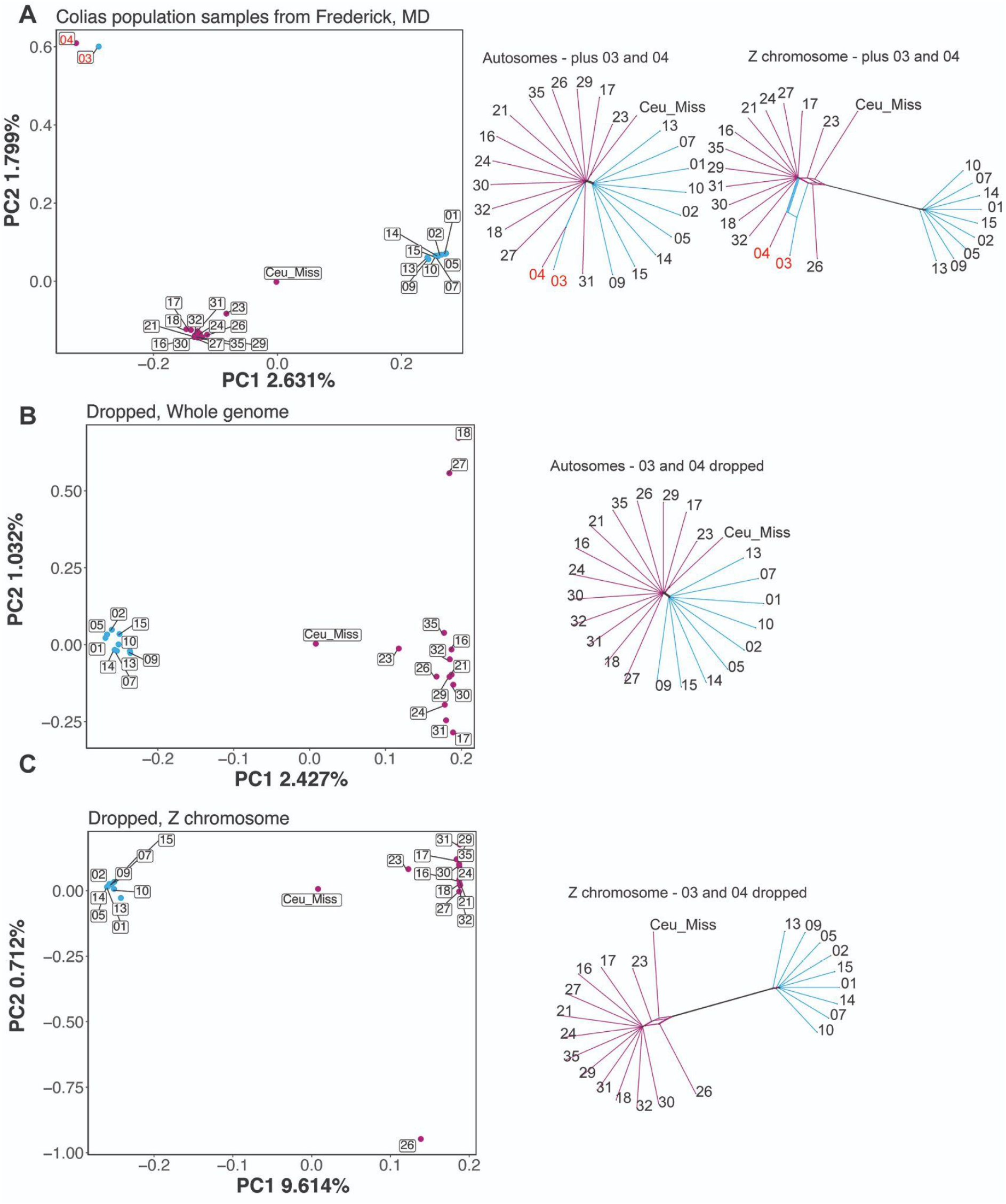
Principal component and phylogenetic network analyses of individuals collected in central Maryland. **A**. Genome-wide SNP PCA and dendrogram of all males sampled from the admixed Maryland population, plus a *C. eurytheme* male collected in Mississippi. **B**. As in A, but omitting the two samples that did not cluster with either species (shown in red in panel A).**C**. Analysis repeated for just the Z chromosome. Individual genomes were assigned to a *C. eurytheme* (magenta) or *C. philodice* (cyan) cluster in subsequent population scans.

**Fig. S3.**
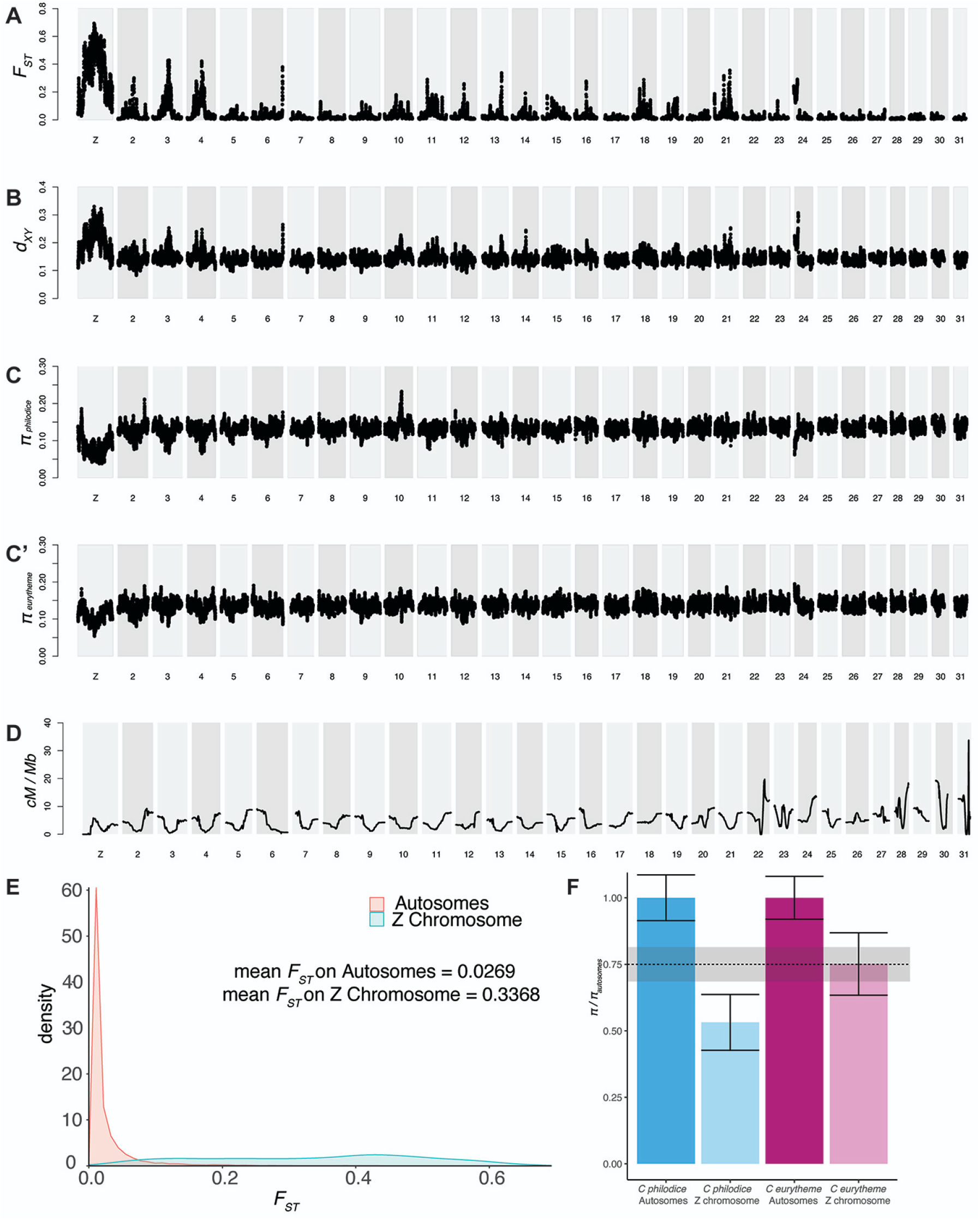
Population statistics for the central Maryland population of *C. philodice* and *C. eurytheme*. **A**. Unadorned *F*_*ST*_ Manhattan plot replicated from figure 1, (**B**) *d*_*XY*_, and (**C-C’**) *π* statistics for each species. **D**. Genome-wide recombination rates derived from the linkage map from the backcross and F2 broods (**Fig. S5**). Ends of chromosomes tend to show higher recombination than centers, and shorter chromosomes were found to have a higher recombination rate than longer ones. This inverse relationship between chromosome size and recombination rate is reminiscent of previous findings in *Heliconius* butterflies (75). **E**. Histogram of *F*_*ST*_ on autosomes versus the Z chromosome. **F**. *π*_*A*_*/π*_*A*_ vs. *π*_*Z*_*/π*_*A*_, with the dotted line and shading representing the expected mean value and confidence interval for the *π*_*Z*_*/π*_*A*_ ratio around 0.75.

**Fig. S4.**
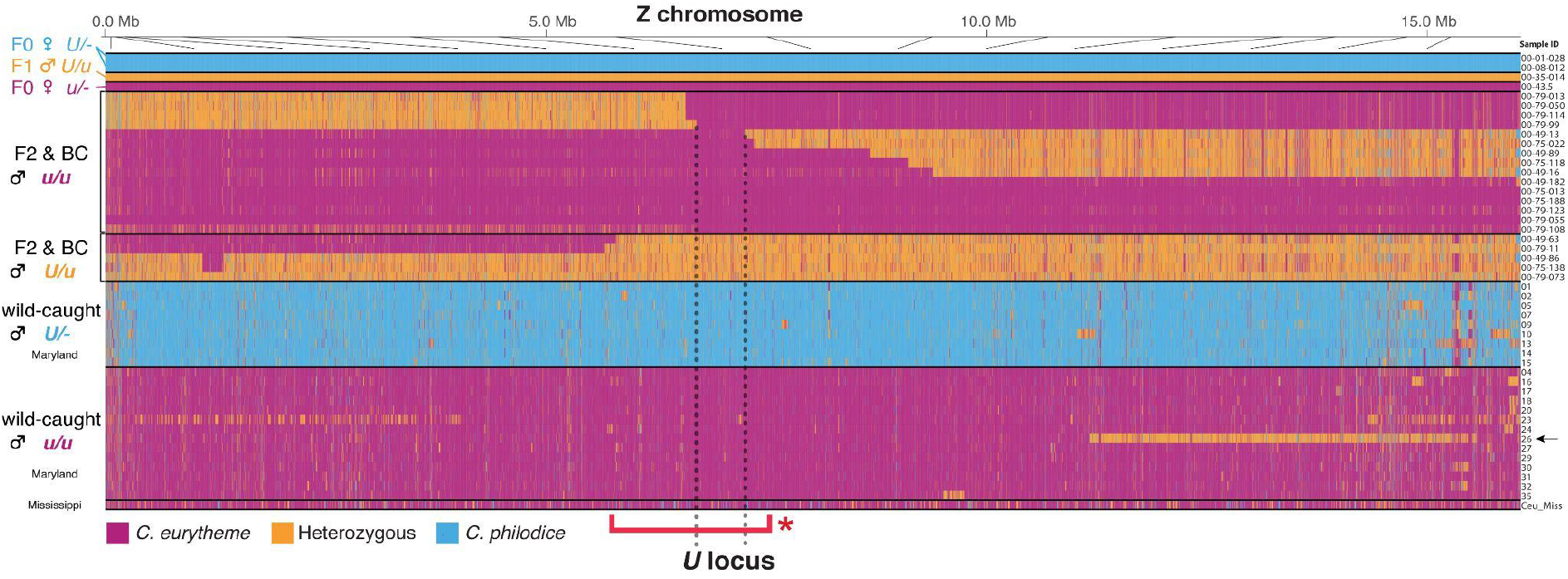
Genotype plot of the Z chromosome for all sequenced samples. A reproduction of **Fig. 1F**, including the samples collected in central Maryland and the *C. eurytheme* individual from Mississippi. The dotted lines and red bracket indicate the same intervals as in **Fig. 1F**. Sample *26* (arrow) has a heterozygous haplotype that takes up a large portion of the Z chromosome, and other individuals of both *C. eurytheme* and *C. philodice* carry shorter heterozygous haplotypes. This provides evidence that some limited ongoing sex-chromosome admixture exists in this wild population, permitting some recombination and persistance in the heterozygous state.

**Fig. S5.**
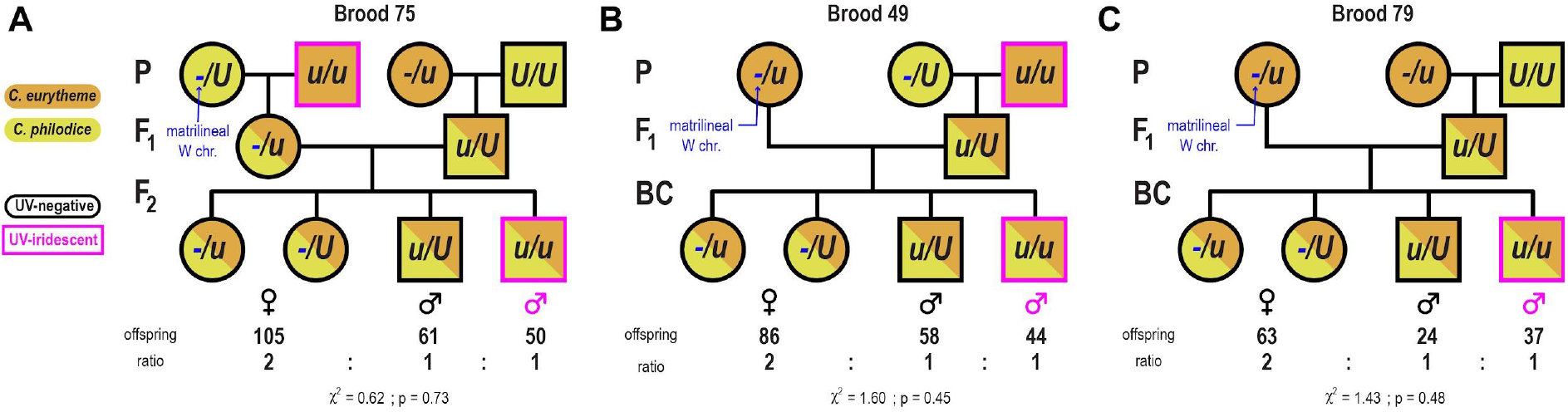
Mapping crosses. The UV trait segregated at the expected 1:1 ratio among the male offspring, in the three crosses that were used for fine mapping the *U* locus. **A**. F2 cross. **B-C**. Backcrosses to *C. eurytheme*, with the second *u* allele inherited from the paternal grandfather (**B**) or paternal grandmother (**C**). Chi-square tests show non-significant deviations from 2:1:1 distributions (females : non-UV males : UV males) that are expected from a Mendelian recessive, sex-linked mode of inheritance of the UV-iridescent state.

**Fig. S6.**
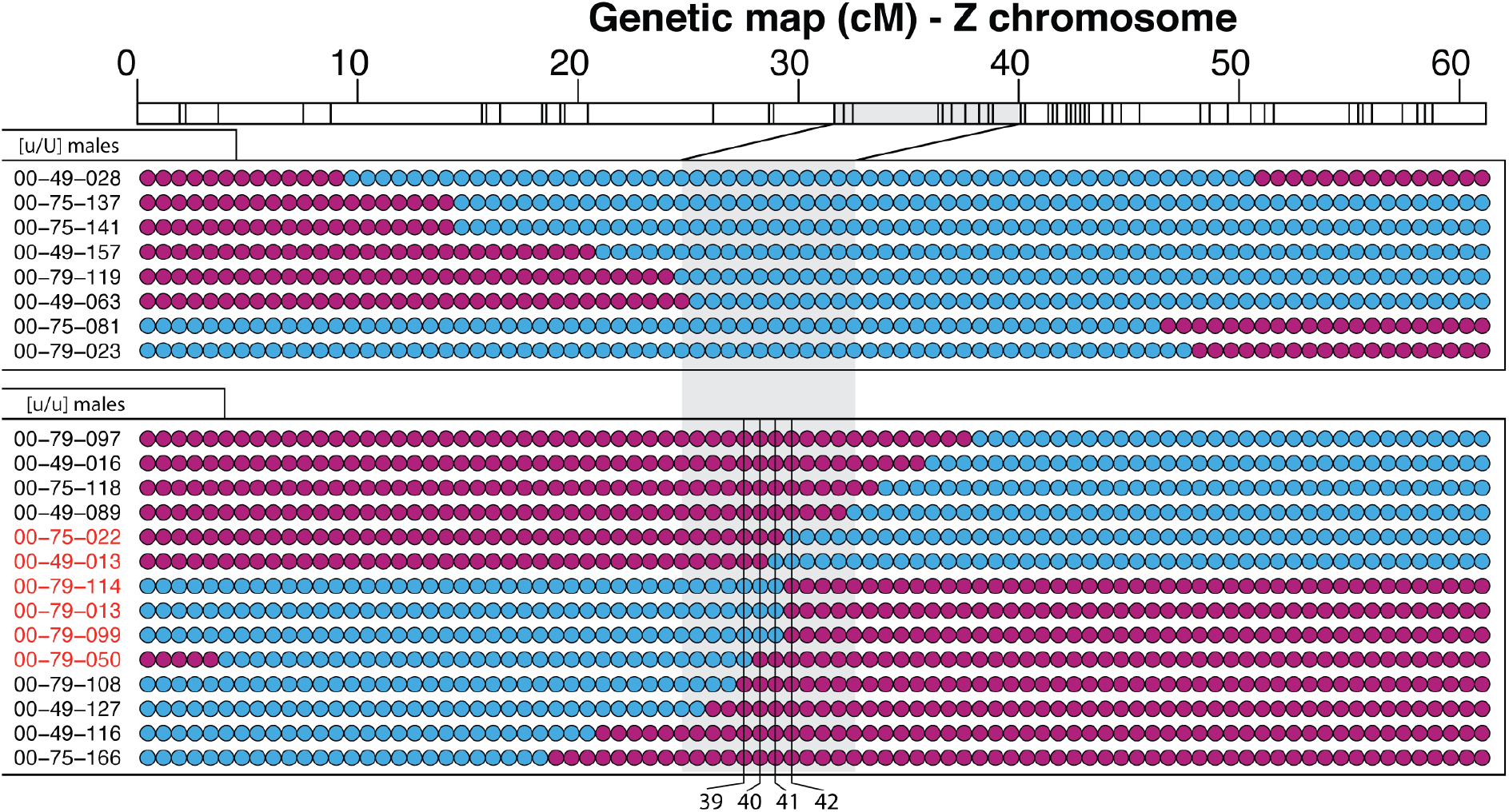
Identification of recombinant individuals in 2b-RAD data. Each row is a recombinant, patrilineal Z chromosome from a male individual, each column is a 2b-RAD marker. All F2 and BC male offspring have a non-recombining matrilineal Z chromosome of *C. eurytheme* ancestry (*i*.*e*. bearing an *u* allele), due to the nature of the crosses (**Fig. S5**). Colors indicate allelic states across the recombining, patrilineal Z chromosome — *magenta* : *C. eurytheme* ancestry ; *blue* : *C. philodice* ancestry. The scale at the top shows the linkage map for the Z chromosome, with the 1.5-LOD support interval for the *U* locus shaded in grey. Red-highlighted individuals had recombination breakpoints between markers 39-42 and were used (along with additional samples) for whole genome resequencing.

**Fig. S7.**
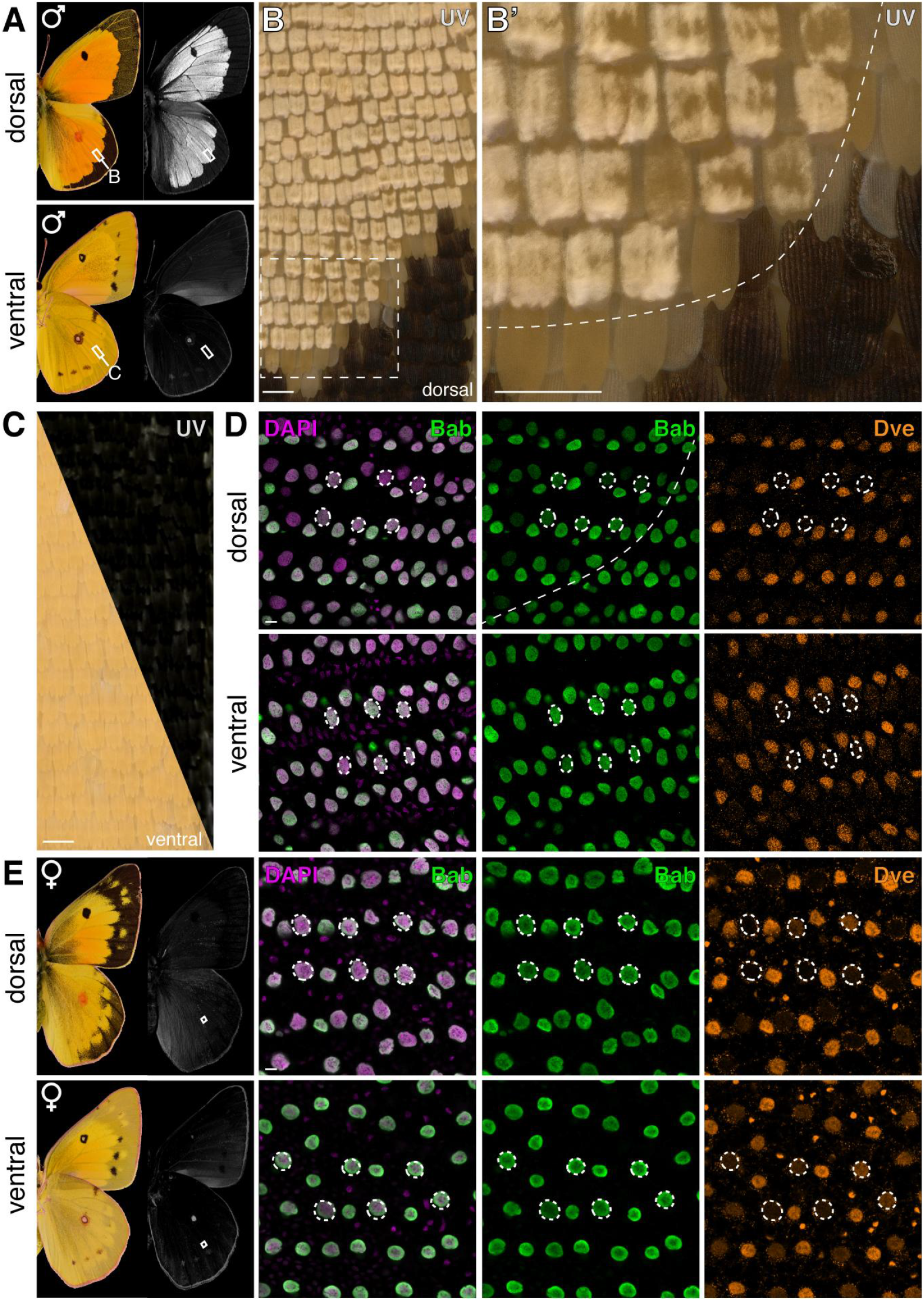
Bab expression is restricted to non-UV scale cell precursors during *C. eurytheme* pupal wings. **A-C**., UV-photography (white) shows the UV-iridescent cover scales are specific to large orange-yellow areas of the dorsal surface of *C. eurytheme* males. Melanic cover scales, and a fraction of yellow-orange cover scales (*e*.*g*. in **B-B’** at the interface with the melanic margin) are UV-negative. Ventral discal spots show broad-spectrum metallic reflectance rather than UV-iridescence (76). **D**. Immunodetection of Bab in a male pupal wing (46% development), in the region shown in (**B’**). Bab is ubiquitous in the UV-negative wing regions. In the dorsal UV-iridescent region, Bab is limited to Dve-positive ground scales, and is repressed in cover scales (circles). **E**., Ubiquitous expression of Bab and Dve observed in both dorsal and ventral surfaces of a female *C. eurytheme* hindwing (37% development). Scale bars: B-C = 100 μm ; D-E = 10 μm.

**Fig. S8.**
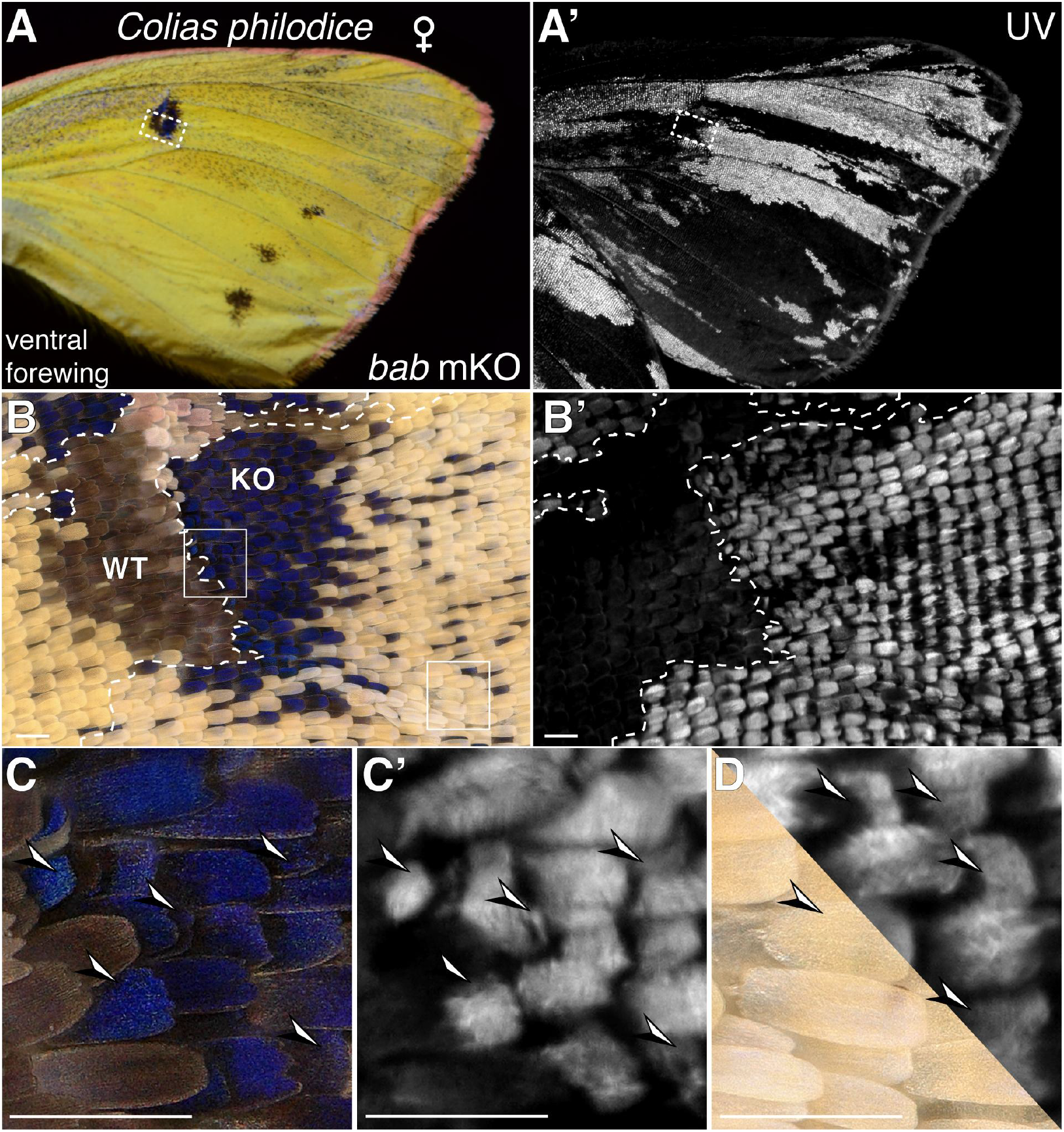
*bab* CRISPR mosaic KOs transform scales to the UV-iridescent state. **A-B**. *bab* mKOs result in gain of UV-iridescence in both pterin and melanin pigment scales, here in the discal spot region of a female *C. philodice* ventral wing. **C-D**. Magnified views from (**A-B**) showing the acquisition of UV-iridescence by mutant ground scales (arrowheads). Scale bars: 100 μm.

**Fig. S9.**
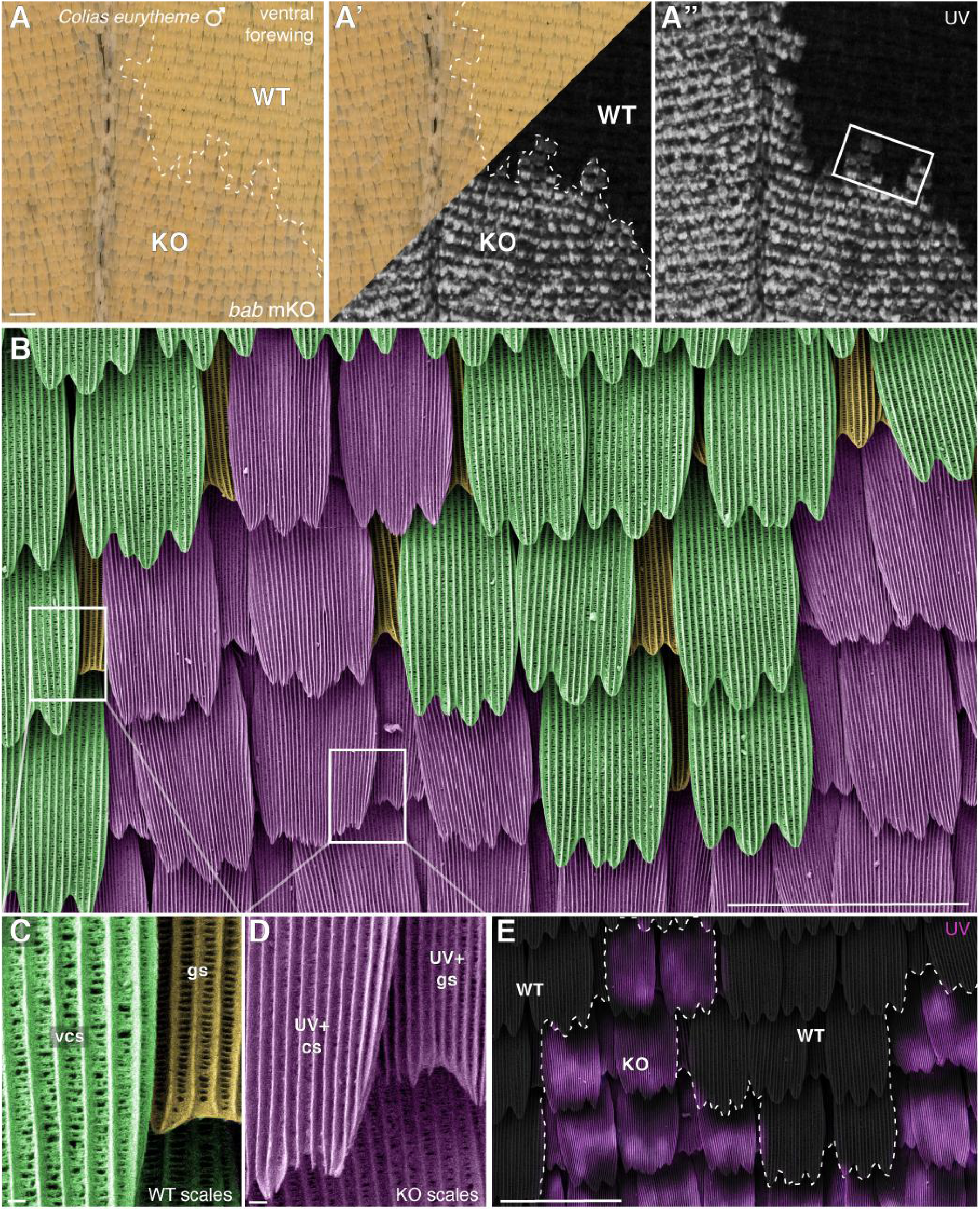
*bab* CRISPR mosaic KOs transform both ground and cover scales to the UV-iridescent state. A-A”. Wild-type ventral forewings of male *C. eurytheme* are normally UV-on the ventral side, but show ectopic UV-iridescence upon mosaic *bab* mKO. **B-D**. False-colored scanning electron micrograph featuring magnified views of both wild-type and *bab* KO clones (B: box in A” ; C-D : boxes in B). Ventral cover scales (green, vcs) and ground scales (gs, orange) are transformed into UV+ scales with dense longitudinal ridges. **E**. Correlated light and electronic microscopy featuring a superimposition of the UV iridescent reflective light onto the SEM view (B). Scale bars: A-B, E = 100 μm ; C-D = 2 μm.

**Fig. S10.**
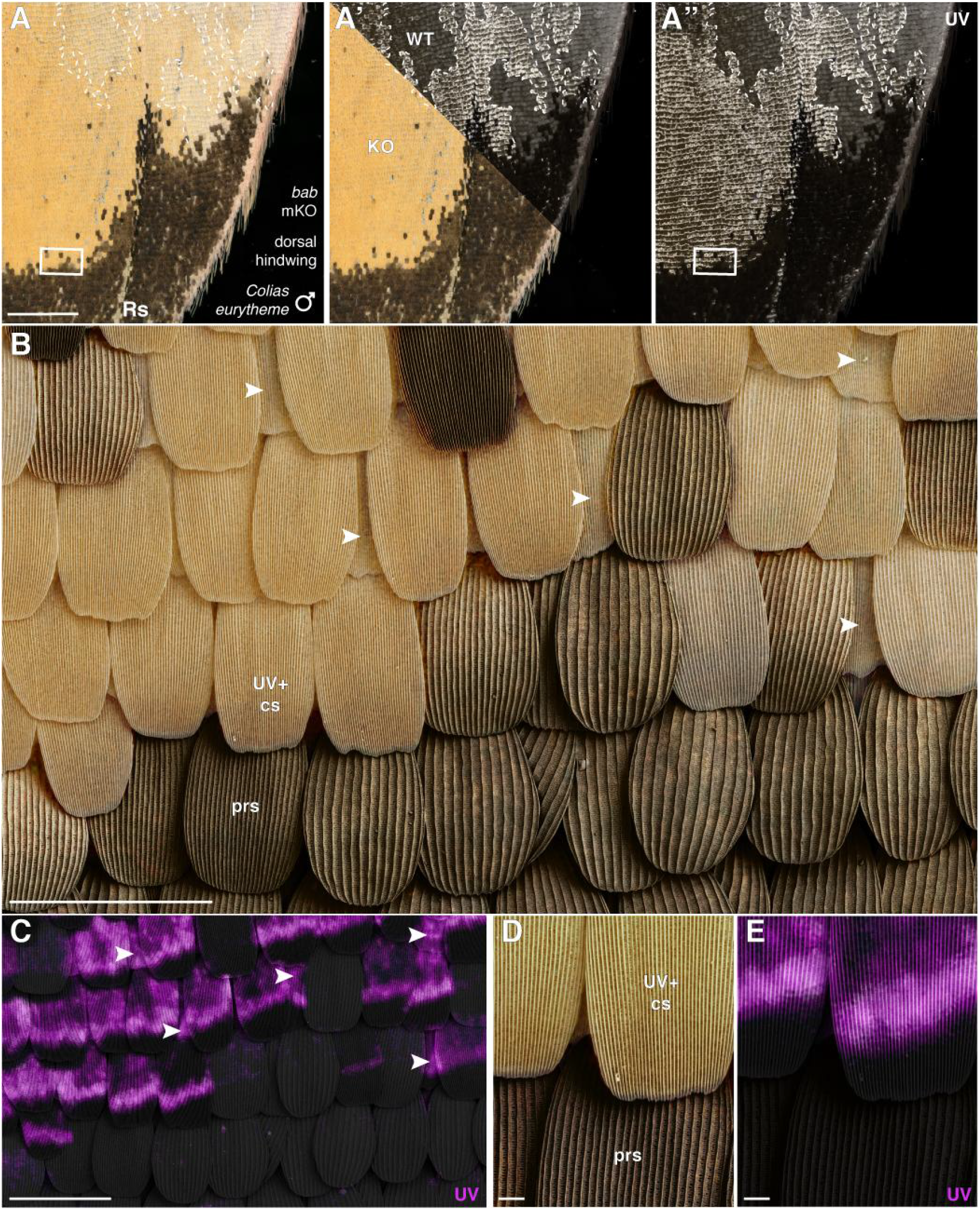
*bab* CRISPR mosaic KOs in dorsal male *C. eurytheme* wings. A-A”. UV-iridescence normally stops posteriorly to the Rs vein in wild-type male *C. eurytheme* dorsal hindwings. In this *bab* mKO wing, UV+ scales extend beyond this boundary, with clones anterior (to the right) of Rs. **B-E**. Correlative light and electron microscope images of a yellow/melanic boundary zone (box in A) featuring an SEM view superimposed with visible (B) or UV-iridescent light (C). The UV-iridescence of ground scales (arrowheads) indicates this area is *bab*-deficient. Unlike other melanic scales from either sex, pheromone-retention scales (prs) that are exclusively found in the dorsal wing margins of *Colias* males, were never transformed following *bab* mKO. Scale bars: A = 500 μm ; B-C = 100 μm ; D-E = 10 μm.

**Fig. S11.**
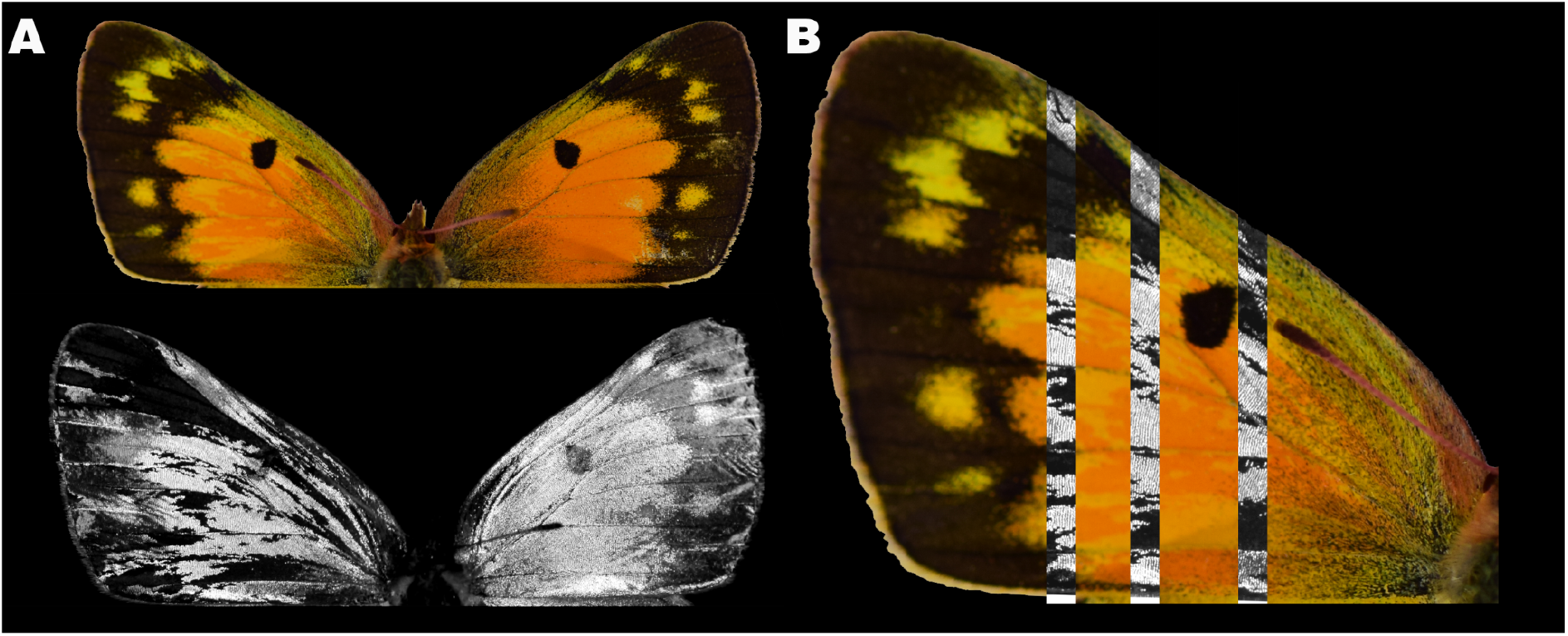
Female-specific effects of *bab* KO on pigmentation. (**A-B**) *C. eurytheme* females exhibit a gain of orange pigmentation in addition to UV transformation. When present, gain of orange is nested in UV transformed clones.

**Fig. S12.**
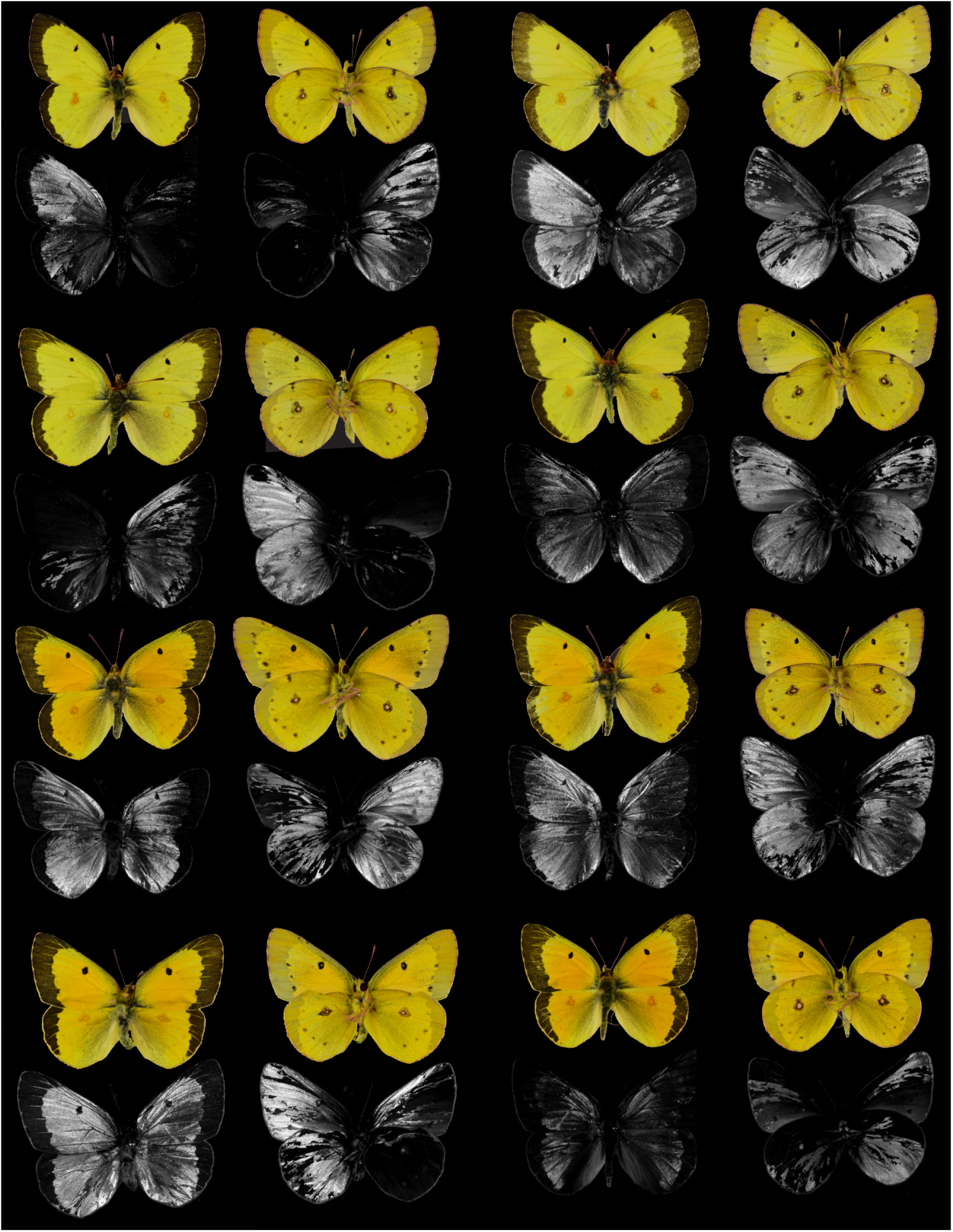
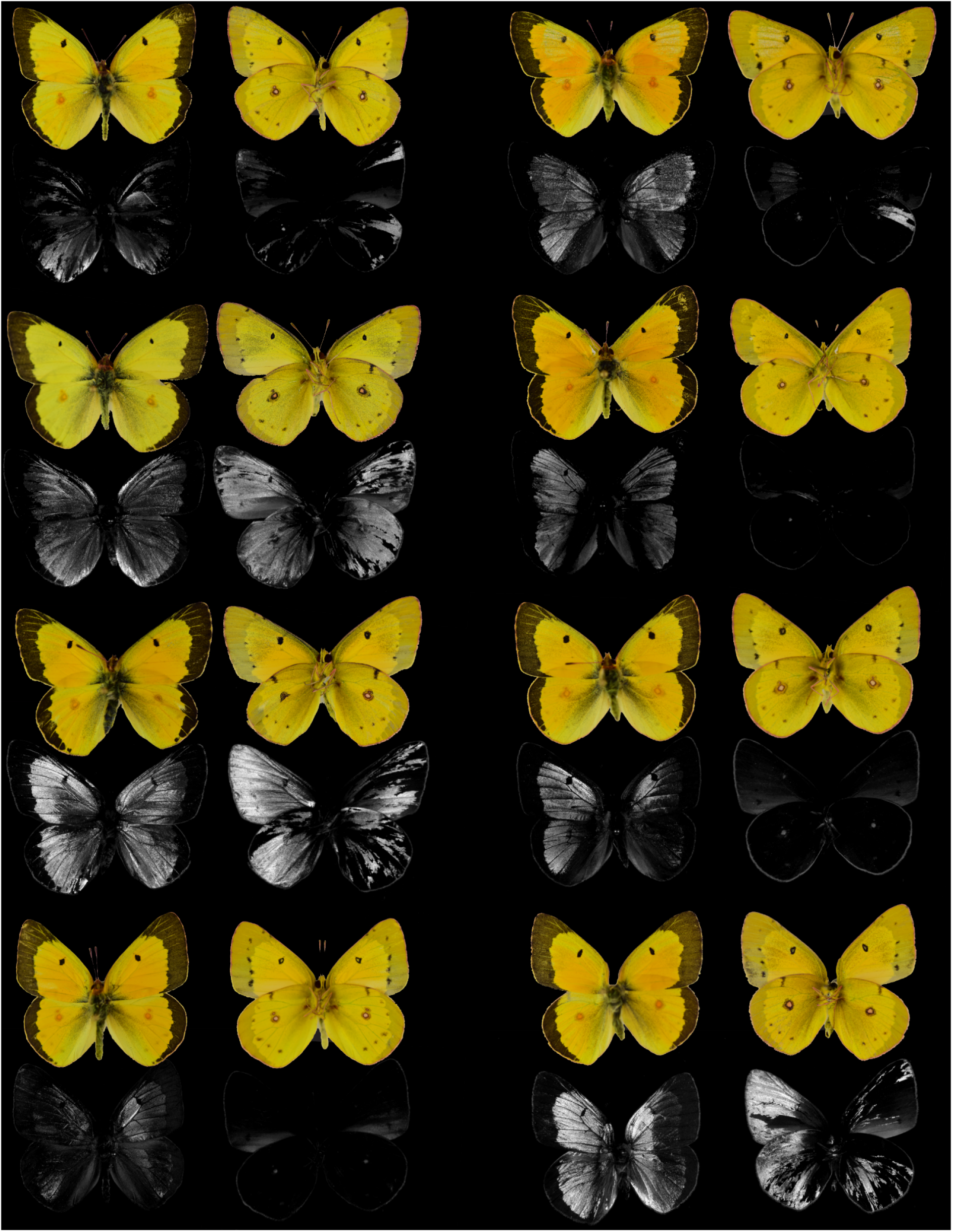
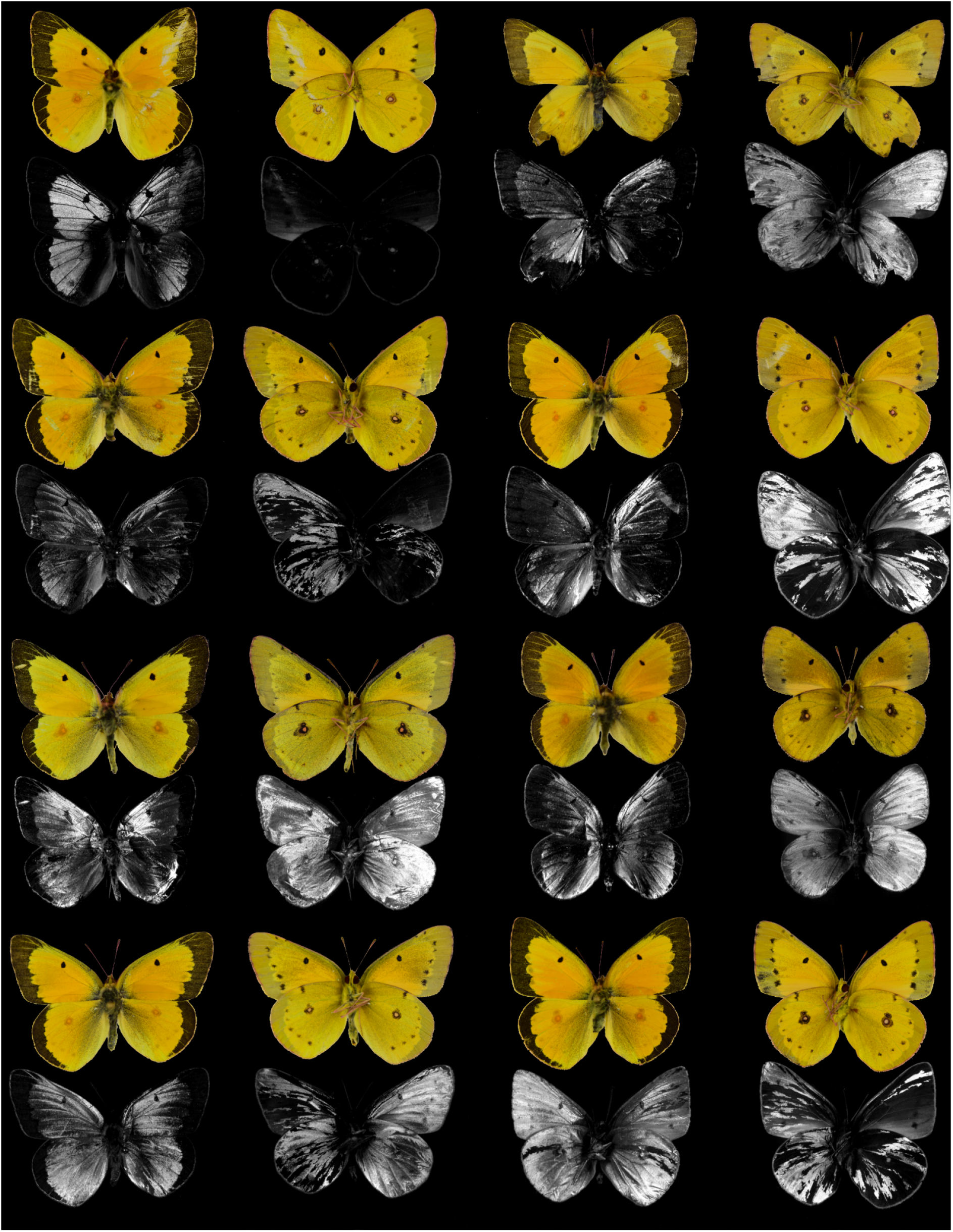
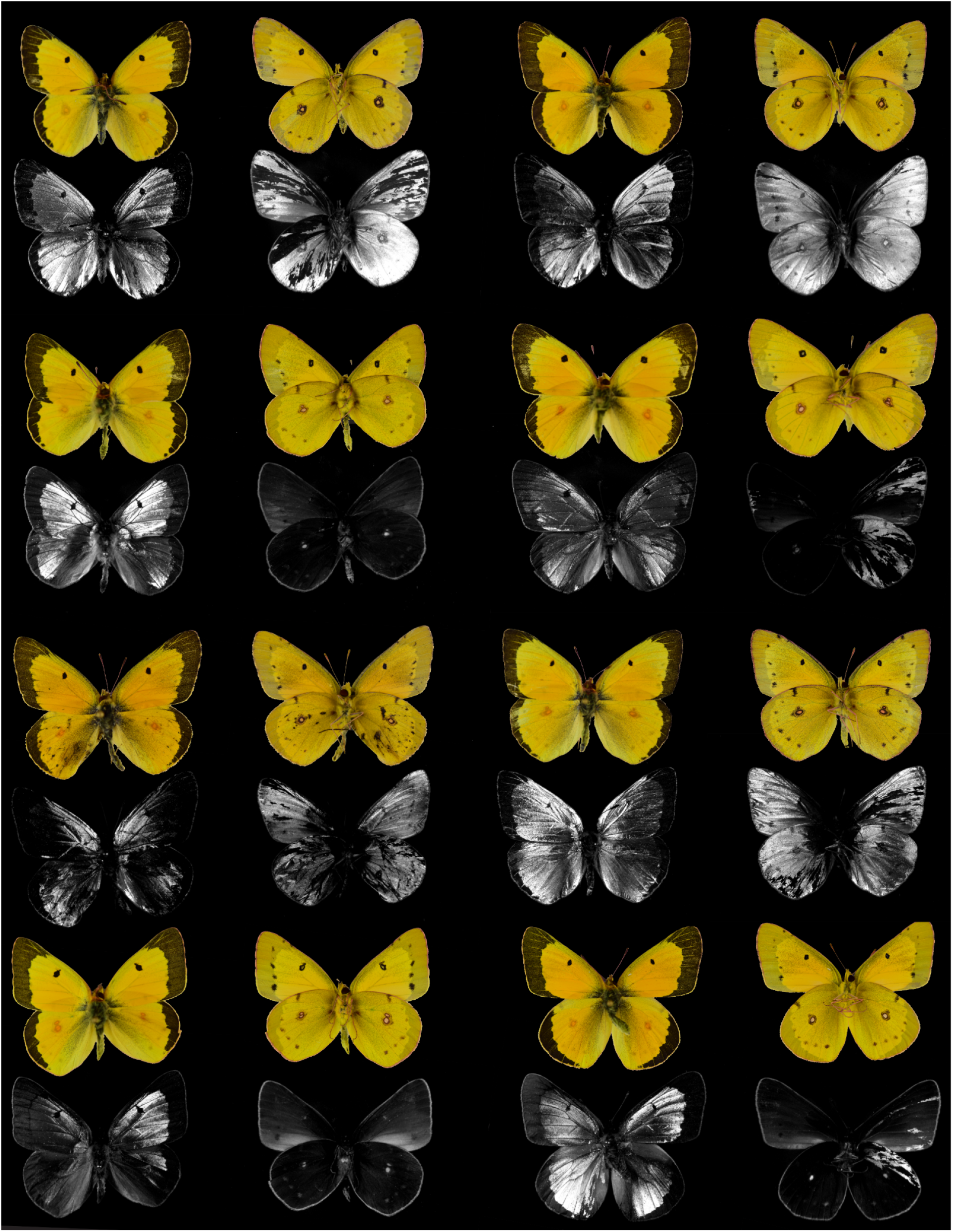
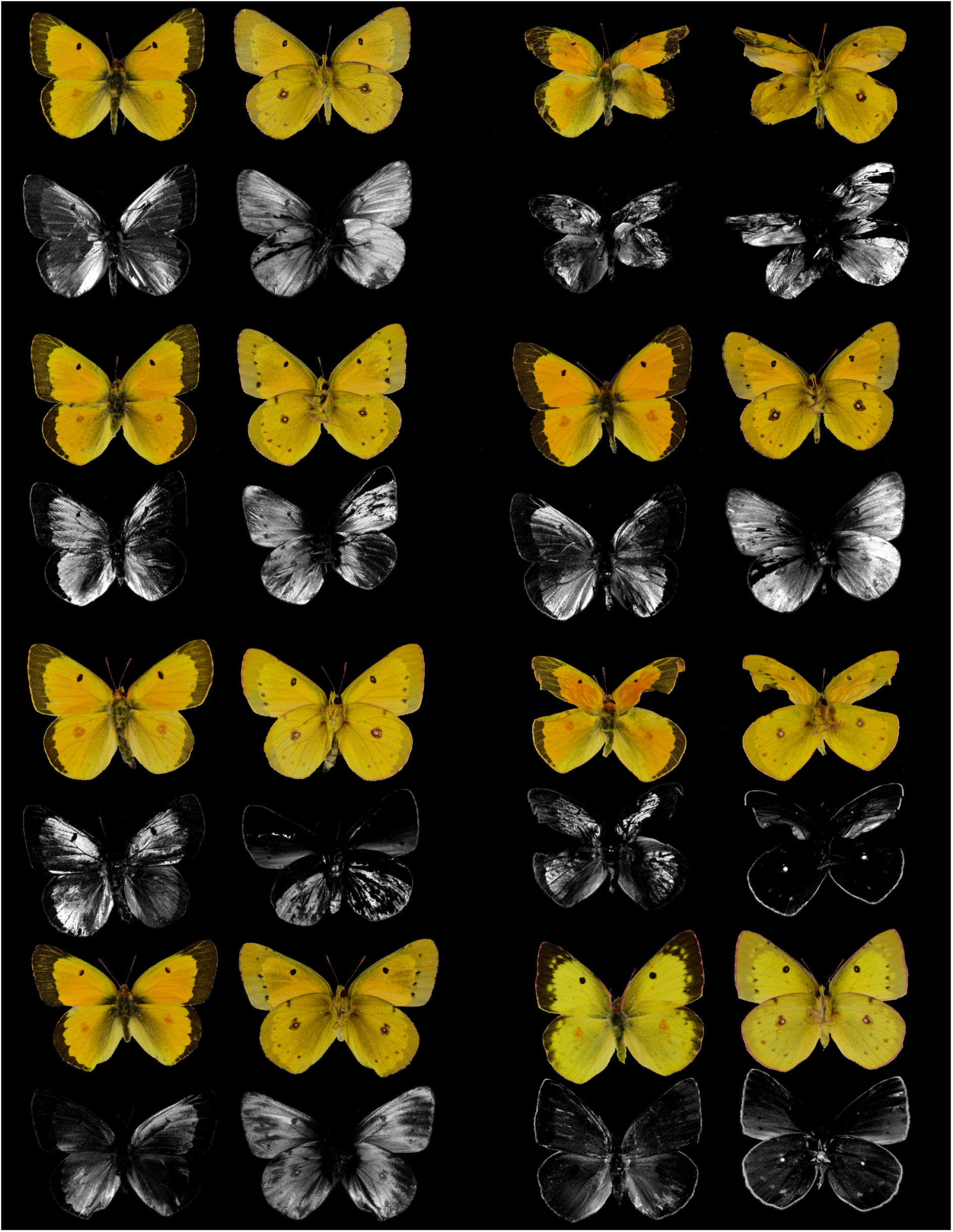
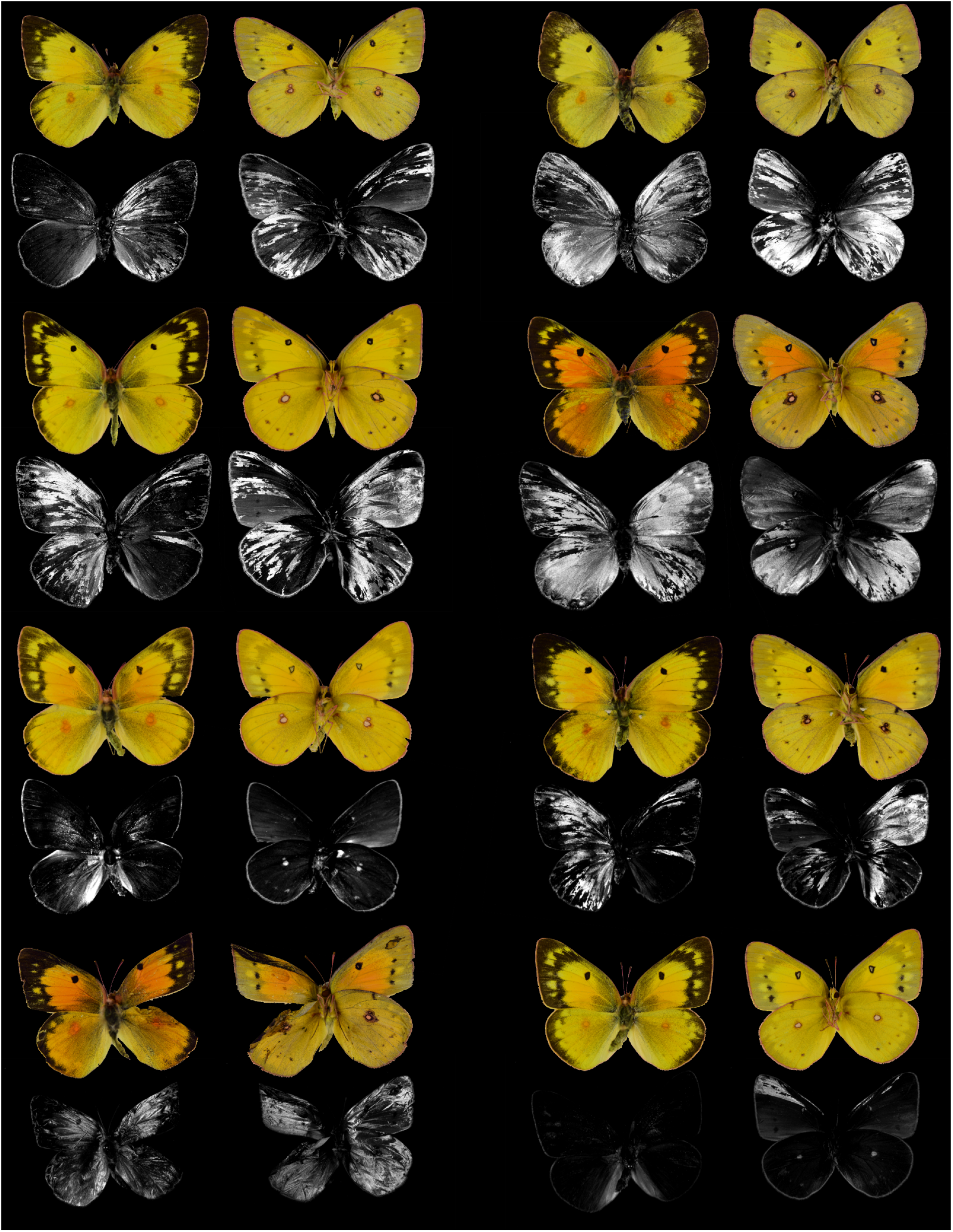
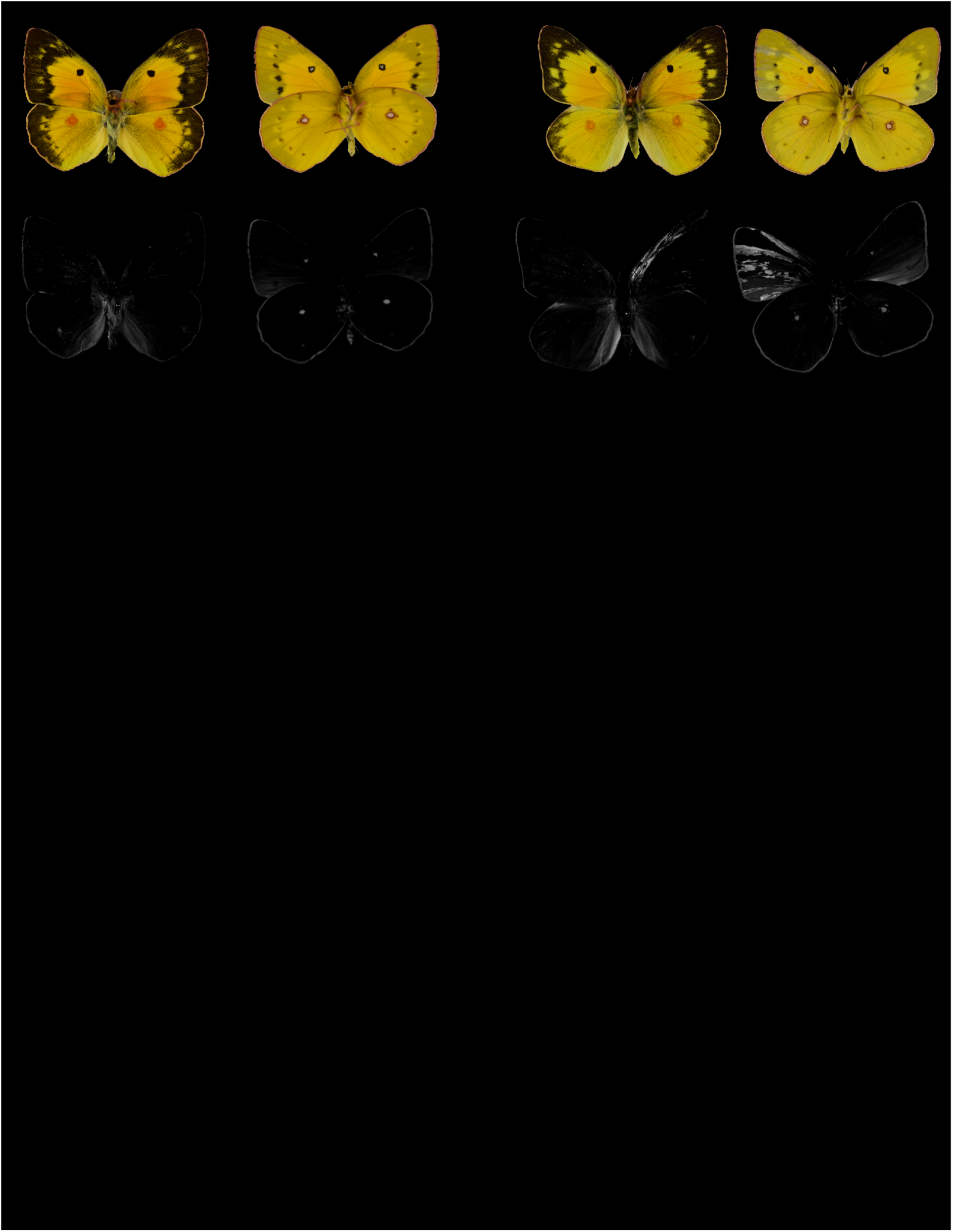
CRISPR G_0_ phenotypes following *bab* mosaic KO. First injection batch.

**Fig. S13.**
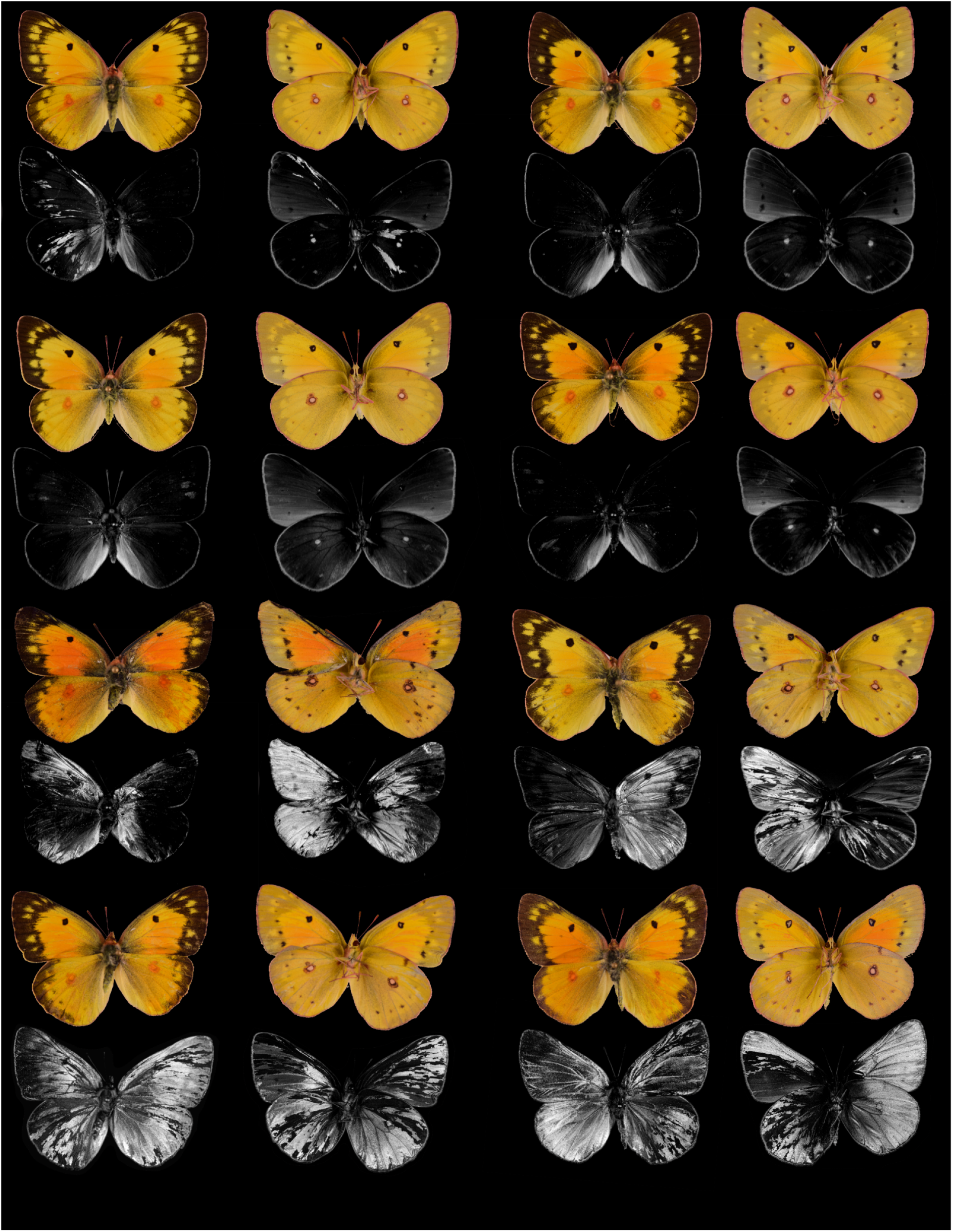
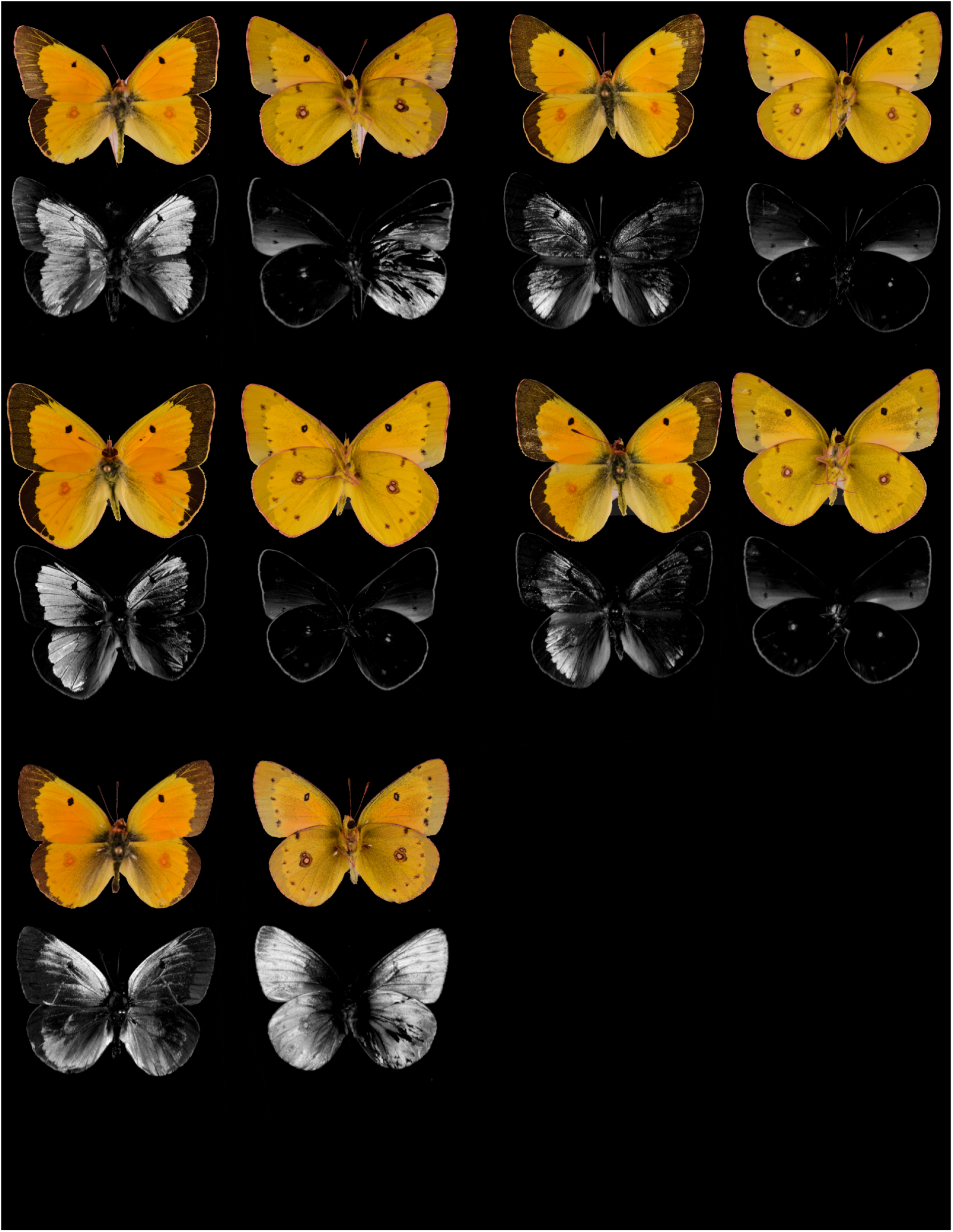
CRISPR G_0_ phenotypes following *bab* mosaic KO. Second injection batch.

## Supplementary Tables

**Table S1.** Resequenced samples for population genomics.

| Individual sample name | SRA Accession Number | Morphospecies | Color | UV trait | Sequencing coverage (x) | Notes |
| --- | --- | --- | --- | --- | --- | --- |
| *Ceu04* | SRR12672378 | *C. eurytheme* | Orange | UV+ | 14.3 | Excluded from genome scan |
| *Ceu16* | SRR12672392 | *C. eurytheme* | Orange | UV+ | 19.4 |  |
| *Ceu17* | SRR12672391 | *C. eurytheme* | Orange | UV+ | 7.4 |  |
| *Ceu18* | SRR12672390 | *C. eurytheme* | Orange | UV+ | 10.2 |  |
| *Ceu21* | SRR12672389 | *C. eurytheme* | Orange | UV+ | 22.1 |  |
| *Ceu23* | SRR12672388 | *C. eurytheme* | Orange | UV+ | 23.2 |  |
| *Ceu24* | SRR12672387 | *C. eurytheme* | Orange | UV+ | 13.1 |  |
| *Ceu26* | SRR12672386 | *C. eurytheme* x *philodice* hybrid | Orange | UV+ | 17.3 | *C. eurytheme* PCA cluster, hybrid Z chromosome |
| *Ceu27* | SRR12672385 | *C. eurytheme* | Orange | UV+ | 15.2 |  |
| *Ceu29* | SRR12672384 | *C. eurytheme* | Orange | UV+ | 10.1 |  |
| *Ceu30* | SRR12672382 | *C. eurytheme* | Orange | UV+ | 12.0 |  |
| *Ceu31* | SRR12672381 | *C. eurytheme* | Orange | UV+ | 10.1 |  |
| *Ceu32* | SRR12672380 | *C. eurytheme* | Orange | UV+ | 15.0 |  |
| *Ceu35* | SRR12672379 | *C. eurytheme* | Orange | UV+ | 18.4 |  |
| *Ceu_Miss* | SRR14305386 | *C. eurytheme* | Orange | NA | 19.8 | Sample from Mississippi |
| *Cph01* | SRR12672395 | *C. philodice* | Yellow | UV- | 18.0 |  |
| *Cph02* | SRR12672394 | *C. philodice* | Yellow | UV- | 13.0 |  |
| *Cph03* | SRR12672383 | *C. philodice* | Yellow | UV- | 8.2 | Excluded from genome scan |
| *Cph05* | SRR12672377 | *C. philodice* | Yellow | UV- | 15.4 |  |
| *Cph07* | SRR12672376 | *C. philodice* | Yellow | UV- | 13.4 |  |
| *Cph09* | SRR12672375 | *C. philodice* | Yellow | UV- | 14.8 |  |
| *Cph11* | SRR12672374 | *C. philodice* | Yellow | UV- | 14.8 |  |
| *Cph13* | SRR12672373 | *C. philodice* | Yellow | UV- | 8.6 |  |
| *Cph14* | SRR12672372 | *C. philodice* | Yellow | UV- | 14.5 |  |
| *Cph15* | SRR12672393 | *C. philodice* | Yellow | UV- | 14.6 |  |

**Table S2.** Summary statistics from the whole-genome population resequencing data.

| **Statistic** | **$d_{XY}$** | **$F_{ST}$** | **$\pi_{philodice}$** | **$\pi_{eurytheme}$** |
| --- | --- | --- | --- | --- |
| Whole genome | 0.146 | 0.042 | 0.130 | 0.136 |
| Autosomes | 0.143 | 0.027 | 0.133 | 0.137 |
| Z-chromosome | 0.205 | 0.337 | 0.071 | 0.103 |

**Table S3.**
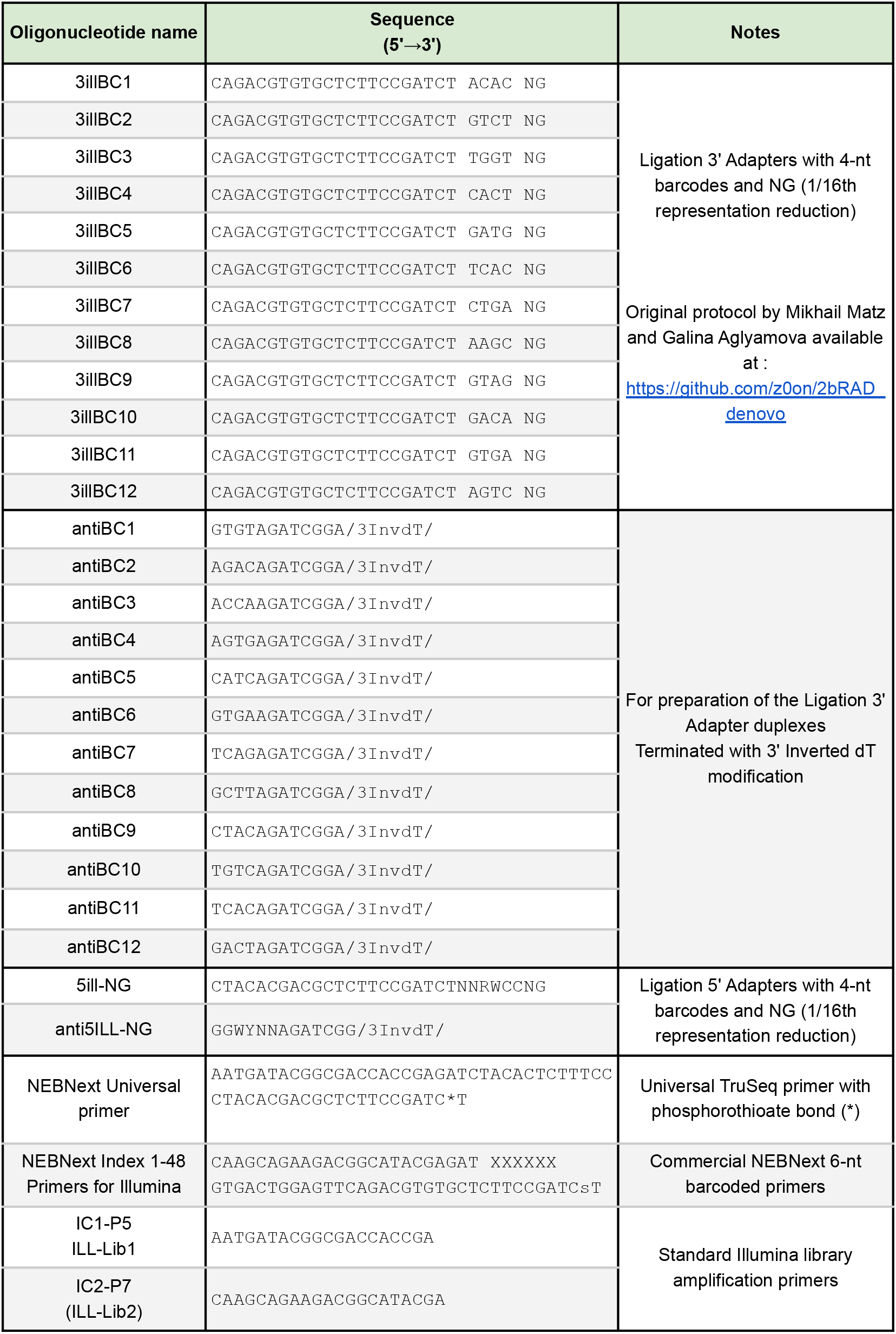
List of oligonucleotides used for 2b-RAD genotyping.

**Table S4.** Resequenced samples from the mapping broods.

| Brood and sample name | SRA Accession Number | Sex | UV trait | Sequencing coverage (x) | Notes |
| --- | --- | --- | --- | --- | --- |
| 00-01-028 | SRR14168767 | f | UV- | 16.7 | *C. philodice*, paternal grandmother of 79 |
| 00-08-012 | SRR14168766 | f | UV- | 19.9 | *C. philodice*, maternal grandmother of 75 |
| 00-35-014 | SRR14168748 | m | UV- | 16.4 | F1, paternal uncle of 75, 49 |
| 00-43.5 | SRR14168747 | f | UV- | 20.2 | *C. eurytheme*, paternal grandmother of 75, 49 |
| 00-49-013 | SRR14168746 | m | UV+ | 16.2 | Recombinant Z : 2b-RAD region 39-42 (Fig. S6) |
| 00-49-016 | SRR14168745 | m | UV+ | 18.5 | Recombinant Z (Fig. S6) |
| 00-49-063 | SRR14168743 | m | UV- | 17.5 | Recombinant Z (Fig. S6) |
| 00-49-086 | SRR14168742 | m | UV- | 17.2 | No recombinant Z |
| 00-49-089 | SRR14168765 | m | UV+ | 20.5 | Recombinant Z (Fig. S6) |
| 00-49-182 | SRR14168744 | m | UV+ | 14.8 | No recombinant Z |
| 00-75-013 | SRR14168764 | m | UV+ | 22.8 | No recombinant Z |
| 00-75-022 | SRR14168763 | m | UV+ | 16.6 | Recombinant Z : 2b-RAD region 39-42 (Fig. S6) |
| 00-75-118 | SRR14168762 | m | UV+ | 16.6 | Recombinant Z (Fig. S6) |
| 00-75-138 | SRR14168761 | m | UV- | 16.6 | No recombinant Z |
| 00-75-188 | SRR14168760 | m | UV+ | 25.6 | No recombinant Z |
| 00-79-013 | SRR14168759 | m | UV+ | 15.8 | Recombinant Z : 2b-RAD region 39-42 (Fig. S6) |
| 00-79-050 | SRR14168758 | m | UV+ | 17.1 | Recombinant Z : 2b-RAD region 39-42 (Fig. S6) |
| 00-79-055 | SRR14168757 | f | UV- | 15.2 | No recombinant Z |
| 00-79-062 | SRR14168756 | m | UV+ | 17.0 | No recombinant Z |
| 00-79-073 | SRR14168754 | m | UV- | 14.9 | No recombinant Z |
| 00-79-099 | SRR14168749 | m | UV+ | 18.3 | Recombinant Z : 2b-RAD region 39-42 (Fig. S6) |
| 00-79-108 | SRR14168753 | m | UV+ | 14.4 | Recombinant Z (Fig. S6) |
| 00-79-114 | SRR14168752 | m | UV+ | 25.4 | Recombinant Z : 2b-RAD region 39-42 (Fig. S6) |
| 00-79-119 | SRR14168751 | m | UV- | 17.9 | Recombinant Z (Fig. S6) |
| 00-79-123 | SRR14168750 | m | UV+ | 26.0 | No recombinant Z |

**Table S5.** Annotated genes in the mapped *U* locus interval.

| *Colias eurytheme* v1<br>Assembly Gene<br>ID | Flybase<br>Orthologous<br>Gene Name | Annotation Summary |
| --- | --- | --- |
| *jg6723* | *Knockout* | Axon guidance |
| *jg6722* | *I(1)G0289* | Plexin domain-containing |
| *jg6721* | *Drat* | Effector of ethanol induced apoptosis |
| *jg6719 ; jg6720* | *BtbVII* | BTB-POZ and Homeobox domain-containing transcription factor |
| *jg6718* | *Ptp61F* | Protein tyrosine phosphatase |
| *jg6717* | *ArfGAP1* | GTPase activating protein |
| *jg6716* | *Pex16* | Peroxisome organization, spermatocyte division, and fatty acid catabolism |
| *jg6714* | *CG9289* | Carboxylic ester hydrolase activity (predicted) |
| *jg6715* | *CG30283* | Serine-type endopeptidase activity (predicted) |
| *jg6713* | *CG42390* | GTPase activity (predicted) |
| *jg6712* | *sim* | Transcription factor, neural development, walking behaviour. |
| *jg6709* | *NimC4* | Phagocytosis of apoptotic cells |
| *jg6701 ; jg6704* | *ets65a* | Transcription factor |
| *jg6703* | *CG16789* | ATP-binding (predicted) |
| *jg6702* | *CG42526* | Transcription regulator (predicted) |
| *jg6700* | *Not1* | Poly-(A) ribonuclease, translation inhibition, ovarian follicle cell development |
| *jg6699* | *CNBP* | mRNA-binding, upregulates dMyc in developing wing |
| *jg6695* | ***bab*** | Transcriptional repressor with known roles in the suppression of male-specific features (*see main text*) |

## Legend for Movie S1

Confocal imaging projections in a *C. eurytheme* male forewing at 46% pupal development, featuring the immunofluorescent detection of Bab (green) in all UV-negative precursors. Magenta: DAPI (nuclei) ; orange: Dve ; circles: cover scale nuclei.

